# The V-type H^+^-ATPase is targeted in anti-diuretic hormone control of the Malpighian ‘renal’ tubules

**DOI:** 10.1101/2022.02.13.480270

**Authors:** Farwa Sajadi, María Fernanda Vergara-Martínez, Jean-Paul V. Paluzzi

## Abstract

Like other insects, secretion by mosquito Malpighian tubules (MTs) is driven by the V-type H^+^-ATPase (VA) localized in the apical membrane of principal cells. In *Aedes aegypti*, the anti-diuretic neurohormone CAPA inhibits secretion by MTs stimulated by select diuretic hormones; however, the cellular effectors of this inhibitory signaling cascade remain unclear. Herein, we demonstrate that the VA inhibitor bafilomycin selectively inhibits serotonin (5HT)- and calcitonin-related diuretic hormone (DH_31_)-stimulated secretion. VA activity increases in DH_31_-treated MTs, whereas CAPA abolishes this increase through a NOS/cGMP/PKG signaling pathway. A critical feature of VA activation involves the reversible association of the cytosolic (V_1_) and membrane (V_o_) complexes. Indeed, higher V_1_ protein abundance was found in membrane fractions of DH_31_-treated MTs whereas CAPA significantly decreased V_1_ abundance in membrane fractions while increasing it in cytosolic fractions. V_1_ immunolocalization was observed strictly in the apical membrane of DH_31_ treated MTs whereas immunoreactivity was dispersed following CAPA treatment. VA complexes colocalized apically in female MTs shortly after a blood-meal consistent with the peak and post-peak phases of diuresis. Comparatively, V_1_ immunoreactivity in MTs was more dispersed and did not colocalize with the V_o_ complex in the apical membrane at 3 hours post blood-meal, representing a timepoint after the late phase of diuresis has concluded. Therefore, CAPA inhibition of MTs involves reducing VA activity and promotes complex dissociation hindering secretion. Collectively, these findings reveal a key target in hormone-mediated inhibition of MTs countering diuresis that provides a deeper understanding of this critical physiological process necessary for hydromineral balance.

**Significance Statement:** The V-type H^+^ ATPase (VA), or proton pump, provides the driving force for transepithelial ion and fluid secretion in insect Malpighian tubules (MTs). While studies have shown diuretic stimulation activates various signaling pathways, including cAMP and downstream effectors promoting increased VA activity, our understanding of anti-diuretic signaling and its potential regulation of the VA remains rudimentary. Herein, we show that CAPA neuropeptide acts through the NOS/cGMP/PKG pathway to inhibit DH_31_-stimulated VA activity, supporting the notion that the anti-diuretic regulation is achieved through dissociation of the VA complexes. These results demonstrate a critical role of VA inhibition and trafficking necessary for anti-diuretic signaling and advances our understanding of the complex neuroendocrine control of the MTs in this important human disease-vector mosquito.

## Introduction

Insect post-prandial diuresis is under rigorous control by neuroendocrine factors (1) acting on the Malpighian ‘renal’ tubules (MTs) to regulate primary urine production. In the yellow fever mosquito, *Aedes aegypti,* several diuretics have been identified that regulate urine production including serotonin (5HT), calcitonin-related diuretic hormone (DH_31_), corticotropin-releasing factor-related diuretic hormone (DH_44_) and leucokinin-related (LK) diuretic hormone (2–5). In female *Aedes* mosquitoes, an anti-diuretic peptidergic neurohormone, CAPA, selectively inhibits DH_31_- and 5HT-stimulated secretion of MTs (4, 6, 7). Insect CAPA neuropeptides are produced in the central nervous system and are evolutionarily related to neuromedin U in vertebrates (8). In the fruit fly, *Drosophila melanogaster*, CAPA peptides have been shown to act through a conserved nitridergic signaling pathway to stimulate diuresis by MTs (9, 10); however, a few other studies have alluded to an anti-diuretic role (11, 12). In contrast, in both larval and adult *A. aegypti*, CAPA peptides inhibit fluid secretion through a signaling cascade involving the NOS/cGMP/PKG pathway (6, 7). Despite this, the anti-diuretic signaling mechanism and downstream cellular targets, such as the ion channels and transporters, remain elusive.

In insect MTs, including *A. aegypti*, the bafilomycin-sensitive V-type H^+^ ATPase (VA), also known as the proton pump, functions as an electrogenic pump allowing the transport of protons from the cytoplasm to the tubule lumen, thus generating a cell-negative membrane voltage (13, 14). This membrane voltage can then drive secondary transport processes such as the cation/H^+^ exchanger or anion/H^+^ cotransporter (15, 16). Originally found in vacuolar membranes of animals and plants, the VA has since been found to be essential in cell function in both invertebrates and vertebrates (17). In insects, the VA is densely located in the apical brush border membrane of tubule principal cells (13, 18), which is rich in mitochondria and endoplasmic reticula, fueling the ATP-consuming proton pump (1, 19). Previous studies have shown VA localization within the apical membrane of principal cells along the entire length of the MTs (19), but absent in stellate cells that express relatively higher levels of the P-type Na^+^/K^+^ ATPase (NKA) (19). Due to stronger VA immunoreactivity observed in MTs (19), and greater ATPase activity by electrophysiological assays (13, 14), the VA is categorized as serving mainly, but not exclusively (20), stimulated transport mechanisms, whereas the NKA serves basic cell housekeeping functions when MTs are undergoing low unstimulated rates of secretion (13, 20, 21).

Stimulation of distinct diuretic hormone receptors can activate various signaling pathways, including elevation of cyclic AMP (cAMP) levels which is known to increase VA activity and assembly in insects (22). *In A. aegypti,* DH_31_, identified as the mosquito natriuretic peptide (23), selectively activates transepithelial secretion of Na^+^ in the MTs, using cAMP as a second messenger (1), and upregulating the VA function to stimulate fluid secretion (24). Similarly, 5HT-stimulated diuresis is also thought to be mediated (at least in part) through the cAMP second messenger pathway (25), activating protein kinase A (PKA) to increase the transepithelial voltage of the basolateral membrane in tubule principal cells (1, 26). In contrast, DH_44_ has been shown to initiate diuresis via the paracellular and transcellular pathways, with higher nanomolar concentrations increasing cAMP and Ca^2+^, influencing both paracellular and transcellular transport, and lower nanomolar concentrations acting through the paracellular pathway only, via intracellular Ca^2+^ (27). Thus, due to its predominant role in fluid secretion, the VA could be a likely target for both diuretic and anti-diuretic factors.

Eukaryotic V-ATPases are a multi-subunit protein composed of up to 14 different polypeptides, which form two major structural complexes. The peripheral V_1_ complex (400-600 kDa), is invariably present in the cytoplasm and interacts with ATP, ADP, and inorganic phosphate (28). The cytosolic V_1_ complex consists of eight different subunits (A-H): a globular headpiece with three alternating subunits A and B forming a hexamer with nucleotide binding sites located at their interface, a central rotor stalk with single copies of subunits D and F, and lastly, a peripheral stalk made up of subunits C, E, G, and H (28). The B subunit, shown to have high sequence similarity amongst several species from fungi to mammals (29, 30), is a 56 kDa polypeptide, and is one of the two sites (along with subunit A) (31) in the V_1_ complex that binds ATP. In contrast, the membrane-integrated V_o_ complex (150-350 kDa) mediates the transport of H^+^ across the membrane (28) and is composed of at least six different subunits, which collectively function in the proton translocation pathway (13, 28). Although the proton channel of the VA can be blocked pharmacologically by the macrolide antibiotic, bafilomycin (32), there are two known intrinsic mechanisms for VA regulation: firstly, through oxidation of the cystine residue on the A subunit of the V_1_ complex, thus preventing ATP hydrolysis; secondly, through reversible disassembly of the V_1_ complex from the holoenzyme (13, 33). While the role and regulation of the VA by diuretic hormones in insect MTs has been studied (34–36), research examining anti-diuretic signaling mechanisms involving the VA remains in its infancy.

Considering that the *A. aegypti* mosquito is recognized as a disease vector for several viruses, further investigating the regulatory mechanisms of diuresis and anti-diuresis in the MTs can allow for useful insights into novel methods to control disease transmission. This study aimed to identify the cellular targets necessary for CAPA-mediated inhibition of fluid secretion by MTs stimulated by select diuretic factors in adult female *A. aegypti*. Our results provide evidence that CAPA neuropeptides inhibit fluid secretion in the mosquito by VA complex dissociation, thus hindering VA function and activity that is essential for driving rapid post-prandial diuresis.

## Results

### Bafilomycin inhibits DH_31_- and 5HT-stimulated fluid secretion rate

To determine the appropriate concentration of bafilomycin to test on adult *A. aegypti* MTs, several doses were applied against DH_31_-stimulated tubules (S1A Fig). Higher doses of bafilomycin (10^-4^ M and 10^-5^ M) resulted in significant inhibition of fluid secretion rate, with maximal inhibition leading to a five-fold decrease, observed with treatment of 10^-5^ M bafilomycin. Next, to determine whether inhibiting the VA would decrease fluid secretion rate stimulated by other diuretic hormones, including 5HT and DH_44_, the effect of 10^-5^ M bafilomycin on adult tubules stimulated with these diuretics was tested (**Fig 1**). Fluid secretion rates were measured over 30 min under control (stimulated) conditions, and then at 10-min intervals in the presence of bafilomycin.

**Fig. 1.**
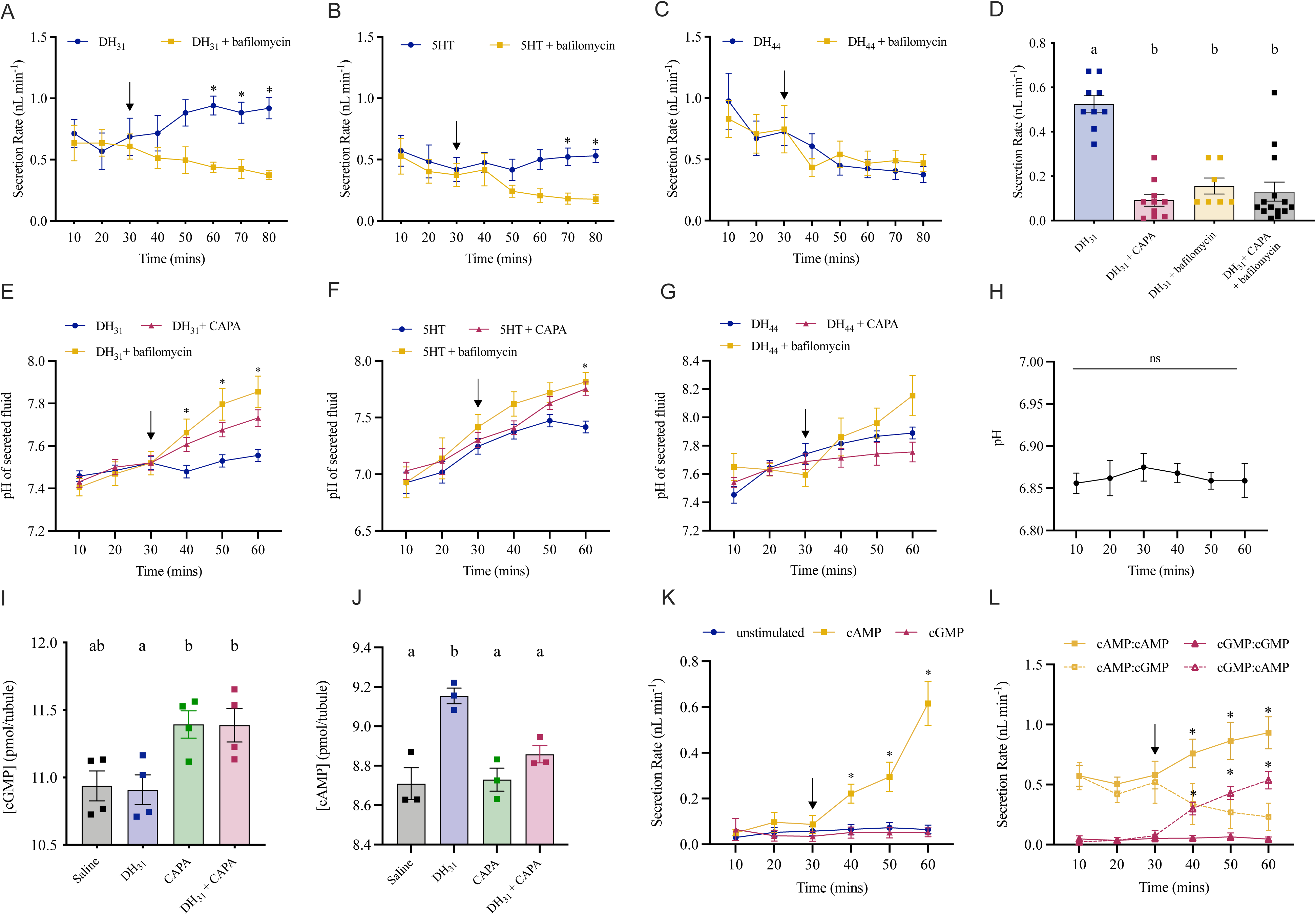
Effect of bafilomycin on fluid secretion rates and pH along with cyclic nucleotide second messengers on adult *A. aegypti* MTs. Tubules were treated with either (A) DH_31_ (B) 5HT or (C) DH_44_ and secreted droplets were measured at 10-min intervals for 30 min. Immediately following measurement of the 30-min point fluid droplet (solid arrow), MTs were treated with (A) DH_31_ (B) 5HT (C) or DH_44_ alone or in combination with *Aedae*CAPA-1 or bafilomycin. (D) MTs were treated with DH_31_ alone or in combination with *Aedae*CAPA-1, bafilomycin, or both *Aedae*CAPA-1 and bafilomycin for 60 min. Secreted fluid pH was measured in tubules treated with either (E) DH_31_ (F) 5HT or (G) DH_44_ before and after addition of *Aedae*CAPA-1 or bafilomycin, along with unstimulated MTs (H). Production of (I) cGMP and (J) cAMP in DH_31_-stimulated MTs treated with *Aedae*CAPA-1. (A-C) Significant differences between bafilomycin-treated and the corresponding time point controls and (E-H) significant differences in secreted fluid pH between *Aedae*CAPA-1- and bafilomycin-treated and the corresponding time point controls are denoted by an asterisk, as determined by a two-way ANOVA and Bonferroni multiple comparison post-hoc test (p<0.05). Data represent the mean ± standard error (n=12-34), ns denotes no statistical significance. (D, I, J) Bars labeled with different letters are significantly different from each other (mean± SEM; one-way ANOVA with Bonferroni multiple comparison, p<0.05, (D) n=7–17) (I, J) n=50 sets of MTs for all treatments (n=3 per treatment)). (K) Fluid secretion rates were measured at 10 min intervals initially over a 30 min interval (unstimulated) and then over a second 30 min interval after the addition (solid arrow) of 10^-4^ M cAMP or 10^-8^ M cGMP. Significant differences between cAMP-treated MTs and the corresponding time point controls (or cGMP-treated MTs) are denoted by an asterisk (mean± SEM; two-way ANOVA with Bonferroni multiple comparison, p<0.05, n=5–6). (L) Significant differences between cAMP-treated MTs and corresponding time point after addition at 30 min (downward arrow) of cGMP are denoted by an asterisk (similar with cGMP alone and cAMP added at 30 min, mean± SEM; one-way ANOVA with Bonferroni multiple comparison, p<0.05, n=5–9).

Treatment of MTs with 10^-5^ M bafilomycin against DH_31_ led to a decrease of fluid secretion over the treatment interval. Specifically, 30 min after treatment with bafilomycin, fluid secretion rate was significantly reduced by over two-fold to 0.438±0.041 nL min^-1^, compared to DH_31_ alone, 0.941±0.077 nL min^-1^ (**Fig 1A**). Similar results were seen with 5HT-stimulated MTs; however, a decrease in fluid secretion was observed 30 min post-bafilomycin, with a significant inhibition 40 min after treatment (0.522±0.072 nL min^-1^, 5HT alone vs. 0.182±0.045 nL min^-1^, 5HT + bafilomycin) (**Fig 1B**). Distinct from DH_31_ and 5HT-stimulated tubules, DH_44_-stimulated secretion was insensitive to bafilomycin treatment (**Fig 1C**). To confirm whether *Aedae*CAPA-1 anti-diuresis is mediated by VA inhibition, adult female MTs were treated with either DH_31_ alone or in combination with *Aedae*CAPA-1, bafilomycin, or both (**Fig 1D**). MTs treated with either *Aedae*CAPA-1 or bafilomycin resulted in a significant inhibition of DH_31_-stimulated secretion, and similar inhibition was observed when both *Aedae*CAPA-1 and bafilomycin were applied together with no evidence of any additive inhibitory effects (**Fig 1D**).

### *Aedae*CAPA-1 and bafilomycin alkalinizes secreted fluid in DH_31_- and 5HT-stimulated MTs

The VA pumps protons from the cell into the tubule lumen thus generating an electromotive potential (20, 31) and providing energy to drive the secretion of cations via Na^+^/H^+^ and/or K^+^/H^+^ antiporters (13). An indirect way to measure whether *Aedae*CAPA-1 and bafilomycin inhibits VA activity involves measuring the pH of the secreted fluid from diuretic-stimulated MTs treated with *Aedae*CAPA-1 or bafilomycin (**Fig 1E–1G**). In DH_31_-stimulated MTs treated with *Aedae*CAPA-1, there was an immediate significantly higher pH in the secreted fluid (7.479±0.030) at 40 min relative to control, increasing up to 7.73±0.038 at 60 min (**Fig 1E**). Similarly, pH levels in DH_31_-stimulated MTs treated with bafilomycin were significantly higher (7.66±0.064) relative to control at 40 min, increasing up to 7.855±0.074 at 60 min. Comparable to DH_31_, addition of *Aedae*CAPA-1 or bafilomycin, significantly increased the pH of secreted fluid from 5HT-stimulated MTs to 7.75±0.061 and 7.82±0.083 respectively, at the 60 min mark (**Fig 1F**). In contrast, unlike the effects observed with DH_31_- and 5HT-stimulated MTs, *Aedae*CAPA-1 or bafilomycin did not alter the pH of the secreted fluid in DH_44_-stimulated MTs (**Fig 1G**). The pH increased from 7.4 to 7.9 during the 30-min DH_44_ incubation; however, pH did not change following the addition of *Aedae*CAPA-1 or bafilomycin. Separately, we conducted measurements in unstimulated tubules to verify pH in these small droplets did not drift over a time frame consistent with our above experiments. Unstimulated MTs were allowed to secrete, and the droplets were isolated and their pH was measured over the course of 60 min. Over this incubation period, no change was observed in the pH of secreted droplets from unstimulated MTs (**Fig 1H**), upholding the notion that the alkalinization of secreted fluid observed following *Aedae*CAPA-1 (or bafilomycin) treatment of DH_31_- and 5HT-stimulated MTs is a result of VA inhibition. Additionally, unstimulated MTs treated with either *Aedae*CAPA-1 or bafilomycin resulted in no significant changes in either secretion rate (**S1B Fig**) or pH (**S1C Fig**).

### *Aedae*CAPA-1 increases cGMP and decreases cAMP levels in DH_31_-treated MTs

To further clarify the CAPA signaling pathway involving the second messengers, cGMP and cAMP, we sought to determine changes in levels of these cyclic nucleotides in MTs incubated in DH_31_ alone or combined with *Aedae*CAPA-1. Treatment of MTs with DH_31_ alone had basal levels of cGMP, 10.91±0.109 pmol^-1^/tubule, comparable to saline treated MTs (**Fig 1I**). Treatment of MTs with *Aedae*CAPA-1 resulted in a significant increase in cGMP levels compared to DH_31_-incubated tubules, increasing to 11.39±0.101 pmol^-1^/tubule. Similar results were observed with MTs treated with both DH_31_ + *Aedae*CAPA-1, with significantly increased cGMP levels of 11.39±0.123 pmol^-1^/tubule (**Fig 1I**) compared to MTs treated with DH_31_ alone. In contrast, treatment of MTs with DH_31_ alone led to significantly higher levels of cAMP, 9.153±0.039 pmol^-1^/tubule, while baseline levels of this second messenger were observed in saline, DH_31_ + *Aedae*CAPA-1, and *Aedae*CAPA-1 treated tubules (**Fig 1J**). To further confirm the stimulatory role of cAMP and inhibitory role of cGMP, tubules were treated with either cyclic nucleotide alone and secretion rates were measured (**Fig 1K**). Unstimulated fluid secretion rates were measured over the first 30 min, and then at 10-min intervals with either cAMP or cGMP. Treatment of MTs with 10^-4^ M cAMP led to a significant increase over the treatment interval, with fluid secretion rates increasing to 0.615±0.096 nL min^-1^ at 60 min, compared to 10^-8^ M cGMP (0.052±0.091 nL min^-1^) and unstimulated (0.065±0.091 nL min^-1^) (**Fig 1K**). Finally, to establish whether these cyclic nucleotide second messengers elicit antagonistic control of the MTs in adult *A. aegypti*, tubules were treated initially with cAMP over the first 30 min and then cGMP was added in the presence of cAMP for a subsequent 30 min (**Fig 1L**). Similarly, we also tested the opposite treatment regime where MTs were treated initially with cGMP and subsequently with cAMP added along with cGMP. Treatment of cAMP-stimulated MTs with 10^-8^ M cGMP led to a significant decrease (∼4-fold) over the treatment interval, with secretion rates decreasing to 0.231±0.113 nL min^-1^ at 60 min, compared to 10^-4^ M cAMP alone (0.931±0.134 nL min^-1^). In contrast, cGMP-incubated tubules treated with 10^-4^ M cAMP led to a significant increase (∼10-fold) in secretion rate (0.537±0.072 nL min^-1^) compared to MTs treated with 10^-8^ M cGMP alone (0.045±0.018 nL min^-1^) (**Fig 1L**).

Parallel studies examining cAMP levels were measured in DH_44_-incubated MTs. Treatment of MTs with DH_44_ alone had high levels of cAMP, 9.115±0.0.061 pmol^-1^/tubule, compared to saline treated MTs, 8.709±0.081 pmol^-1^/tubule (**S2A Fig**). Levels of cAMP remained unchanged in tubules treated with both DH_44_ + *Aedae*CAPA-1, 8.954±0.108 pmol^-1^/tubule. To further resolve the cAMP signaling pathway downstream of DH_31_ and DH_44_ stimulated diuresis, a PKA inhibitor (KT5720) was tested against diuretic-stimulated MTs. KT5720 abolished the stimulatory effect of DH_31_, whereas secretion rates by DH_44_-treated MTs remained unchanged (**S2B Fig**).

In *D. melanogaster*, the effects of CAPA peptides is dependent on both intracellular and extracellular sources of Ca^2+^ (9, 37). Thus, we sought to test the involvement of Ca^2+^ in the CAPA intracellular second messenger pathway in the *Aedes* mosquito. The tubules were tested in a modified Ca^2+^-free saline containing L-glutamine, resulting in secretion rates similar to MTs incubated in Ca^2+^-free saline with Schneider’s medium and *Aedes* saline used previously (**S3A Fig**). Treatment of MTs in the presence of the Ca^2+^-chelating agent EGTA had no effect on the inhibition of DH_31_-stimulated secretion by *Aedae*CAPA-1 (**S3B Fig**). Similarly, inhibition of intracellular Ca^2+^ levels with TMB-8 or membrane permeable BAPTA-AM alone or in combination with EGTA had no effect on the anti-diuretic activity of *Aedae*CAPA-1 (**S3B Fig**).

### *Aedae*CAPA-1 decreases VA activity in DH_31_-stimulated MTs through the NOS/cGMP/PKG pathway

In *Aedes* MTs, 50-60% of the total ATPase activity can be attributed to a bafilomycin- and nitrate-sensitive component that reflects the activity of the VA pump (13). The remaining ATPase activity may be due to nucleotide cyclases, protein kinases, myosin, DNA helicases, and other ATP-consuming processes such as the NKA (13). As such, to determine whether CAPA inhibits VA and/or NKA function in the female mosquito, diuretic-stimulated MTs were challenged with *Aedae*CAPA-1 to measure the resultant NKA and VA activity. As expected, given its established role as the natriuretic diuretic hormone, adult female MTs treated with DH_31_ resulted in a significant (over two-fold) increase of VA activity, 0.0329±0.0007 μmoles ADP/μg protein/hour, compared to saline controls, 0.0151±0.0021 μmoles ADP/μg protein/hour (**Fig 2A**). Importantly, MTs incubated with both DH_31_ and *Aedae*CAPA-1 had significantly lower VA activity, resulting in activity levels indistinguishable from saline controls. In contrast, neither 5HT nor DH_44_ influenced VA activity (p>0.05) when compared with saline controls, while co-treatment with *Aedae*CAPA-1 also resulted in indistinguishable VA activity. Similar VA activity levels were observed between 5HT and DH_44_ (0.0255±0.0078 and 0.0208±0.0042 μmoles ADP/μg protein/hour) and with co-application of *Aedae*CAPA-1 (0.0150±0.0036 and 0.0154±0.0070 μmoles ADP/μg protein/hour). Unlike changes observed in VA activity following treatment with DH_31_, diuretic-stimulation or *Aedae*CAPA-1 treatment did not perturb NKA activity, with levels similar to unstimulated MTs (**Fig 2B**).

**Fig 2.**
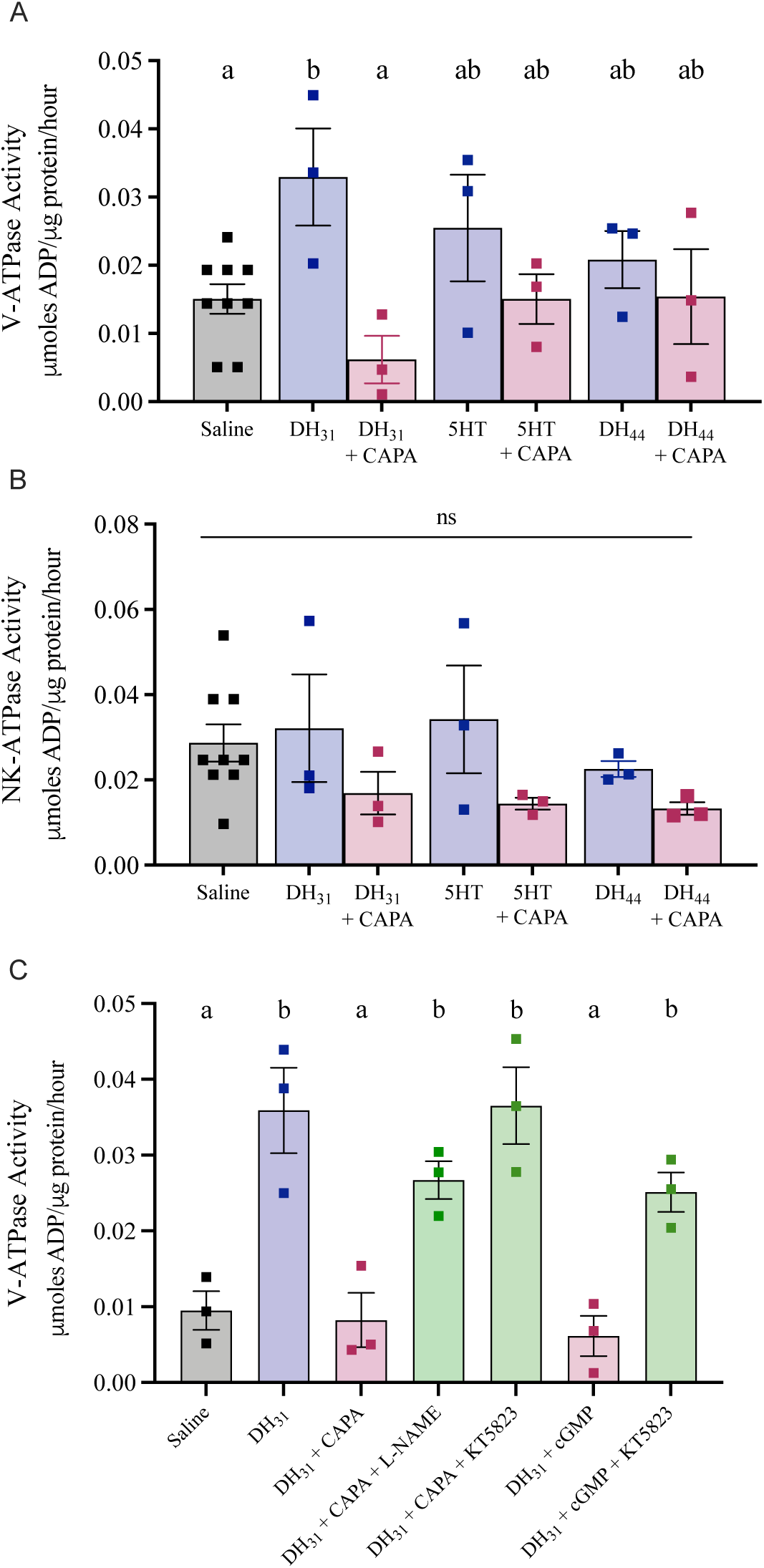
Effect of *Aedae*CAPA-1 and NOS/PKG inhibitors on VA and NKA activity in diuretic stimulated *A. aegypti* MTs. MTs were incubated in *Aedes* saline, diuretics (DH_31_, 5HT, and DH_44_) alone or in combination with *Aedae*CAPA-1 for 30 min before collection to measure (A) VA and (B) NKA activity. (C) MTs were treated with pharmacological blockers, NOS inhibitor (_L_-NAME) and PKG inhibitor (KT5823) in combination with either *Aedae*CAPA-1 or cGMP. Bars labeled with different letters are significantly different from each other (mean± SEM; one-way ANOVA with Bonferroni multiple comparison, p<0.05). N.S. denotes no statistical significance. For each treatment, 50 sets of MTs were collected with n=3 biological replicates per treatment.

To confirm the actions of CAPA are mediated through the NOS/cGMP/PKG pathway, pharmacological blockers, including inhibitors of NOS (_L_-NAME) and PKG (KT5823), were tested against DH_31_-stimulated MTs treated with either *Aedae*CAPA-1 or cGMP (**Fig 2C**). Application of _L_-NAME or KT5823 abolished the inhibitory effect of *Aedae*CAPA-1, resulting in high levels of VA activity, 0.02671±0.0025 and 0.03653±0.0051 μmoles ADP/μg protein/hour respectively, compared to MTs treated with DH_31_ + *Aedae*CAPA-1. As expected, treatment of DH_31_-stimulated MTs with cGMP resulted in a significant decrease in VA activity, 0.006±0.0026 μmoles ADP/μg protein/hour, similar to *Aedae*CAPA-1-treated MTs, while co-treatment with KT5823, abolished the inhibitory effect of cGMP, resulting in an increase in VA activity (**Fig 2C**).

### *Aedae*CAPA-1 leads to VA holoenzyme dissociation in DH_31_-treated MTs

The reversible dissociation of the V_1_ complex from the V_o_ membrane-integrated complex is a well-known mechanism for regulating VA transport activity (22, 31, 34, 35, 38). To determine whether *Aedae*CAPA-1 influences VA complex dissociation, membrane and cytosolic protein fractions were isolated from DH_31_ and DH_31_ + *Aedae*CAPA-1 incubated MTs, and a polyclonal V_1_ antibody (13) was used to measure protein abundance. First, membrane and cytosolic protein isolation was verified with specific membrane (AQP1, **S4A Fig**) and cytosolic (beta-tubulin, **S4B Fig**) markers. Western blot analysis revealed three protein bands, with calculated molecular masses of 74 kDa, 56 kDa, and 32 kDa (13, 39) (**S4C Fig**). The V_1_ complex is composed of eight subunits (A-H), which includes the A (∼74kDa) and B subunit (∼56Da) that are arranged in a ring forming the globular headpiece for ATP binding and hydrolysis (31). Additionally, studies have suggested that subunit D (∼32kDa) alongside subunit F constitute the central rotational stalk of the V_1_ complex (17). There was no difference in abundance observed for the A subunit (74 kDa band) in either membrane or cytosolic fractions between saline and DH_31_ treatments whereas DH_31_ + *Aedae*CAPA-1 incubated MTs had increased A subunit (74 kDa band) protein abundance in cytosolic fractions compared to saline treatment (**Fig 3A**) and decreased abundance in the membrane fraction compared to MTs treated with DH_31_ alone (**Fig 3B**). Similarly, the V_1_ complex B subunit abundance (56 kDa band) was similar in all treatments within the cytosolic protein fraction (**Fig 3C**) whereas DH_31_ + *Aedae*CAPA-1 incubated MTs had significantly lower abundance in membrane fractions compared to MTs treated with DH_31_ alone (**Fig 3D**). Finally, there was no difference in abundance of the V_1_ complex subunit D (32 kDa band) between saline and DH_31_ treated MTs in neither cytosolic nor membrane fractions. However, as observed for the A and B subunit bands (74 and 56 kDa band, respectively) DH_31_ + *Aedae*CAPA-1 incubated MTs showed a significant increase in subunit D abundance in cytosolic fractions compared to saline treated MTs (**Fig 3E**) and a decrease in its abundance in membrane fractions compared to MTs treated with DH_31_ alone (**Fig 3F**). In summary, all three of the V1 complex immunoreactive bands corresponding to subunits A, B and D (74, 56 and 32 kDa, respectively) showed significantly lower abundance in membrane fractions in DH_31_ + *Aedae*CAPA-1 incubated MTs.

**Fig 3.**
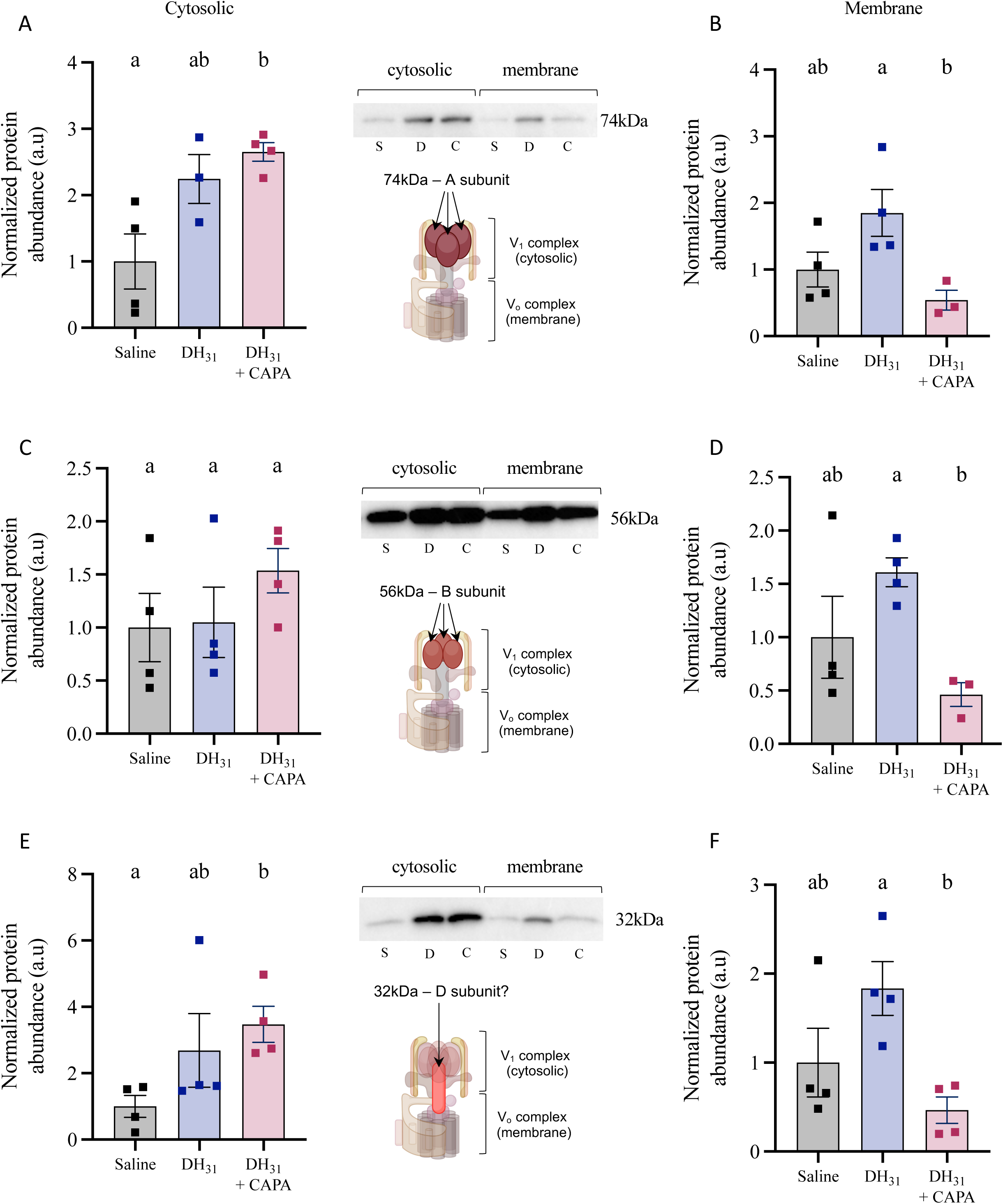
Membrane and cytosolic protein abundance of the V_1_ complex in MTs of *A. aegypti*. The MTs (n=40–50) were incubated in *Aedes* saline, DH_31_, or DH_31_ + *Aedae*CAPA-1 for one hour before collection. Protein abundance was measured in the (A) 74 kDa band, A subunit (B) 56 kDa band, B subunit and (c) 32 kDa band, D subunit of the V_1_ complex. Individual band densities were normalized to total protein using Coomassie staining, and graphed relative to saline-treated controls. Bars labeled with different letters are significantly different from each other (mean± SEM; one-way ANOVA with Bonferroni multiple comparison, p<0.05, n=3–4 replicates).

To visualize this potential endocrine-mediated reorganization of the VA holoenzyme in this simple epithelium, we immunolocalized the membrane-integrated V_o_ and cytosolic V_1_ complex in the female *A. aegypti* MTs. Transverse sections of saline-incubated (control) MTs demonstrated moderate enrichment of V_o_ (**Fig 4A**, **S5A Fig**), and V_1_ (**Fig 4B**, **S5B Fig**) complexes in the apical membrane of principal cells (**Fig 4C**, **S5C–S5D Fig**). Comparatively, DH_31_-incubated MTs revealed intense localization of the V_o_ (**Fig 4D**, **S5E Fig**), and V_1_ (**Fig 4E**, **S5F Fig**) complexes within principal cells, where V_1_ staining was strictly co-localized with V_o_ staining on the apical membrane (**Fig 4F**, **S5G–S5H Fig**). Interestingly, although V_o_ immunolocalization was restricted to the apical membrane (**Fig 4G**, **S5I-K Fig**), V_1_ immunoreactivity was dispersed in both the apical membrane and cytosolic region (**Fig 4H**, **S5L-N Fig**) in DH_31_ + *Aedae*CAPA-1 treated MTs with little evidence of apical co-localization (**Fig 4I**, **S5O-T Fig**) as observed in MTs treated with DH31 alone. Immunostaining was absent in control preparations probed with only secondary antibodies (**S6A-B Fig**) confirming the specific detection of the VA complexes with each primary antibody. Additionally, to further resolve the involvement of the cGMP/PKG pathway in VA holoenzyme organization, DH_31_- and cAMP-stimulated tubules were incubated with cGMP alone or with PKG blocker, KT5823 (**S7 Fig**). As expected, transverse sections of both DH_31_- and cAMP-incubated MTs demonstrated strict colocalization of V_o_ (**S7A, S7J Fig**) and V_1_ (**S7B, S7K Fig**) in the apical membrane of principal cells (**S7C, S7L Fig**), while V_1_ and V_o_ immunoreactivity was found in both the apical membrane and cytosol regions in cAMP + cGMP and DH_31_ + cGMP co-treated MTs (**S7D–F, S7M–O Fig**). However, immunoreactivity of the V_1_ complex was found in the apical membrane alongside the V_o_ complex in preparations treated with the PKG blocker, KT5823 (**S7G–I, S7P–R Fig**).

**Fig 4.**
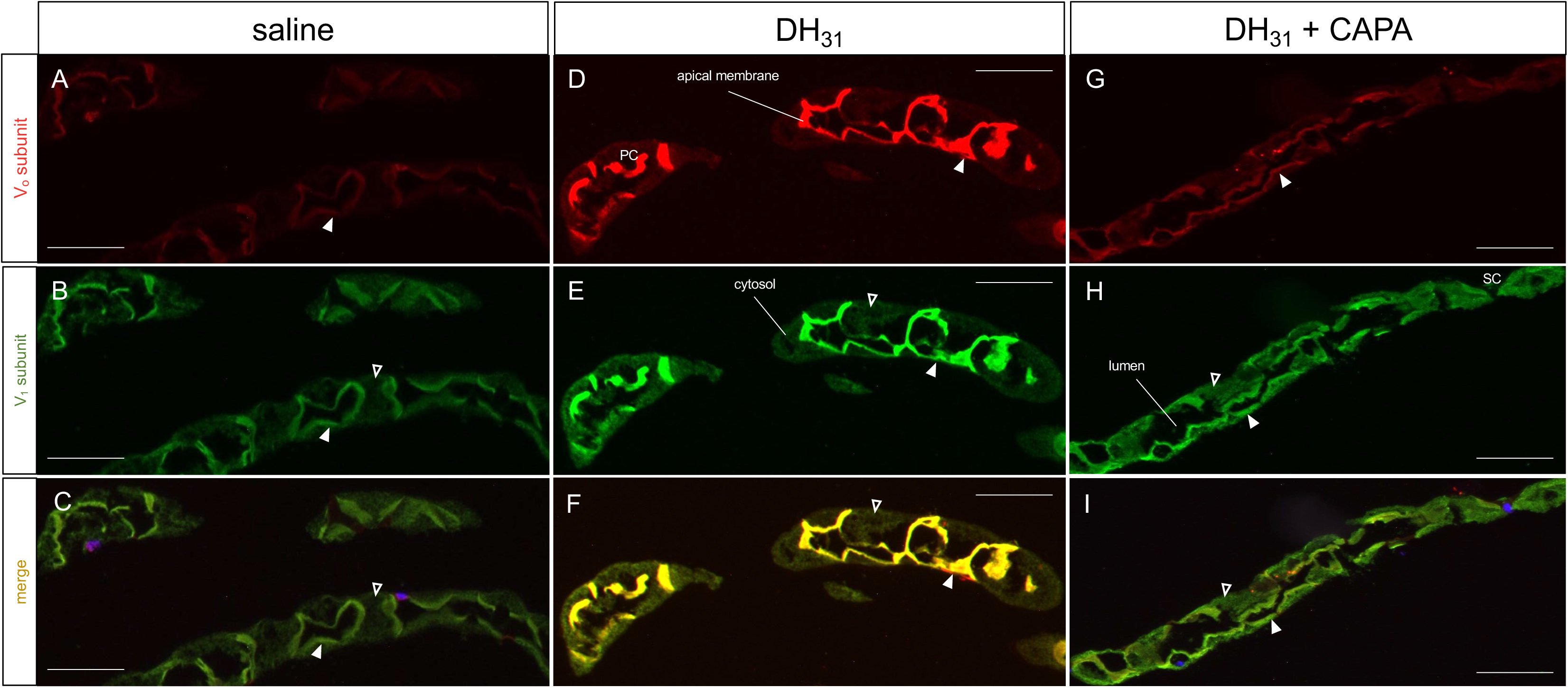
Immunolocalization of the V_o_ and V_1_ complexes in transverse sections of stimulated *A. aegypti* MTs. Representative paraffin-embedded sections of *A. aegypti* MTs incubated in either (A–C) *Aedes* saline alone, (D–F) DH_31_ and (G–I) DH_31_ + *Aedae*CAPA-1 for 30 min. Panels (A,D,G) show V_o_ staining (red), (B,E,H) show V_1_ staining (green), and panels (C,F,I) show merged images with staining highly colocalized in DH_31_ treatment but less evident in saline and *Aedae*CAPA-1 added treatments. Solid white arrows denote apical VA staining, and empty arrows indicate cytosolic VA staining. Where visible in sections, DAPI nuclear staining is shown in blue. Scale bar 100 μm, n = 4 biological replicates (SC = stellate cell).

To investigate this endocrine-mediated phenomenon *in vivo*, we immunolocalized the membrane-integrated V_o_ and cytosolic V_1_ complex in blood-fed females at different time points. Whole body sections of non-blood-fed similarly aged females (control) demonstrated moderate enrichment of both V_o_ (**Fig 5A**), and V_1_ (**Fig 5B**) in the MTs, with minimal co-localization (**Fig 5C**), resembling saline-incubated MTs. Interestingly, blood-fed female MTs revealed strong co-localization of the V_o_ and V_1_ complexes at 10 min (**Fig 5D–5F**) and 30 min (**Fig 5G–5I**) post blood-meal, whereas V_1_ immunoreactivity was more dispersed in both the apical membrane and cytosolic area in MTs 3 hrs post blood-meal (**Fig 5J–5L**), comparable to non-blood-fed females.

**Fig. 5.**
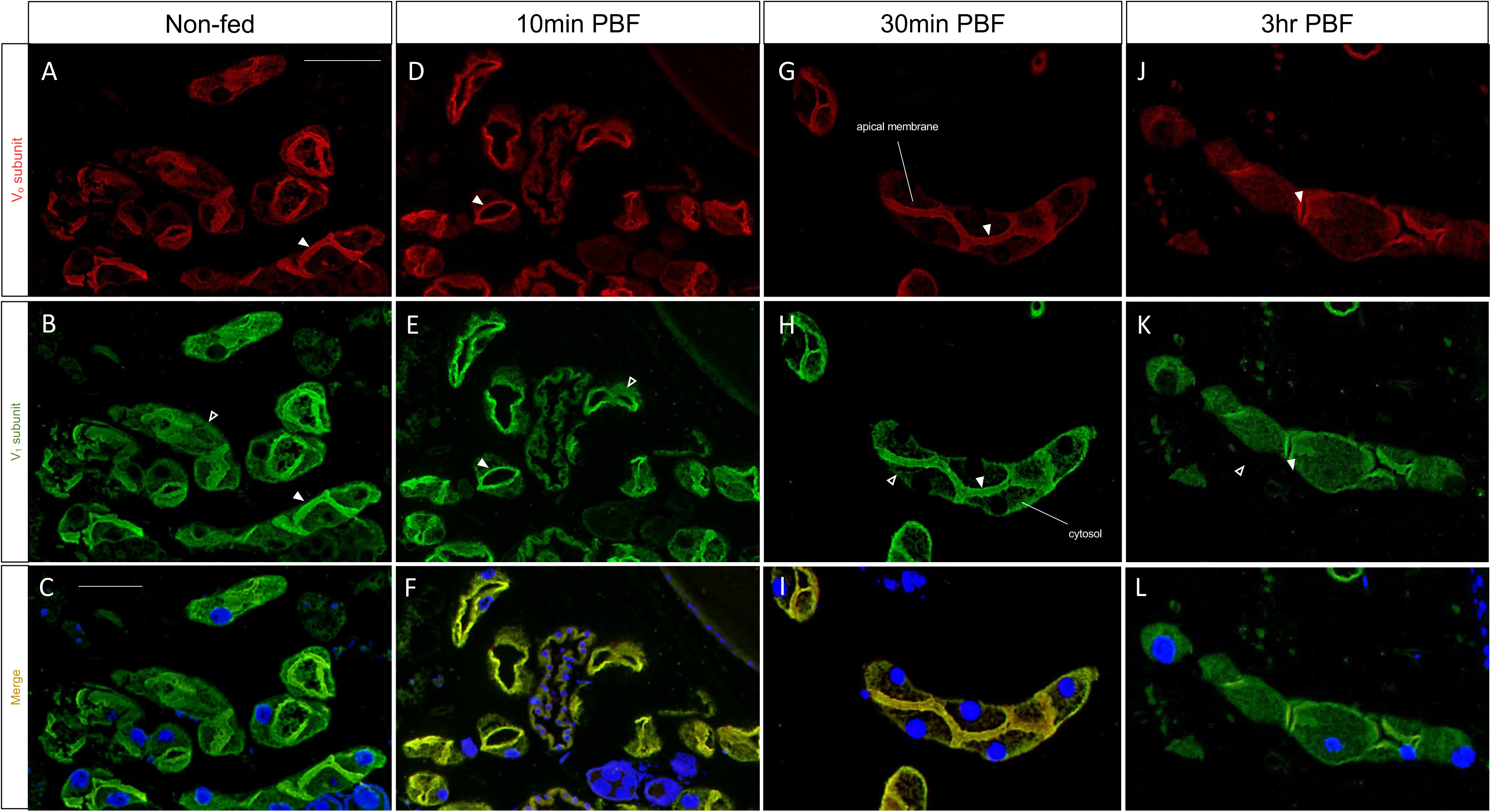
Immunolocalization of the V_o_ and V_1_ complexes in blood-fed *A. aegypti* females. Representative paraffin-embedded sections of whole-body non-blood-fed females (A–C), blood-fed females isolated (D–F) 10 min (G–I) 30 min and (J–L) 3 hr post blood-meal. Panels (A,D,G,J) show V_o_ staining (red), (B,E,H,K) show V_1_ staining (green), and panels (C,F,I,L) show merged immunoreactive staining. DAPI nuclear staining is shown in blue. Scale bar 100 μm, n = 4 biological replicates.

## Discussion

The MTs of the *Aedes* mosquito are the main organs responsible for the secretion of water and solutes, thereby contributing towards hydromineral homeostasis of the animal (40). Active ion transport in *A. aegypti* MTs is accomplished mainly by the V-ATPases (VA) densely localized in the apical brush-border membrane of principal cells, that energize the apical and basolateral membrane as well as the paracellular pathway, allowing for transepithelial secretion of NaCl, KCl, and other solutes (41). In animal cells, VA localized to the plasma membrane, especially on the apical membrane of epithelial cells, contribute to extracellular acidification or alkalization, intracellular pH homeostasis, or energize the plasma membrane for secondary active transport (42, 43). In insect MTs, the VA plays a major role in fluid secretion, thus serving as a primary target for both diuretic and, as this study demonstrates, anti-diuretic hormonal regulation of the mosquito ‘renal’ tubules. Although the structure and function of the VA has been elucidated in some detail (13, 20, 22, 31, 44–46), the regulation of the proton pump remains unclear. Of the various regulatory mechanisms for VA activity, the most studied is the reversible dissociation of the cytosolic V_1_ complex from the membrane-integrated V_o_ complex, first established in the midgut of the tobacco hornworm, *Manduca sexta* and yeast, *Saccharomyces cerevisiae* (45, 47). In this study, the activity and regulation of the VA was investigated under both diuretic and anti-diuretic hormone control of the adult female *A. aegypti* MTs. Notably, the current results advance our knowledge of the anti-diuretic control of the *A. aegypti* MTs, revealing a cellular mechanism for CAPA inhibition of the MTs by targeting the VA to block fluid secretion stimulated by select diuretic factors. This includes inhibition of the DH_31_-related mosquito natriuretic peptide, which is critical for the post-haematophagy diuresis that eliminates excess water and sodium originating from the bloodmeal-derived plasma.

In insects, water excretion is tightly regulated to maintain homeostasis of ions and water (1, 48, 49). Female *A. aegypti* engorge a salt- and water-rich bloodmeal to obtain the necessary nutrients and proteins for their eggs (1), with about 40% of the ingested water eliminated in the first hour post feeding (50). The high rates of water excretion along with the high rates of primary urine production post bloodmeal suggest a highly coordinated and defined hormonal regulation of the signaling processes and downstream cellular targets for ion and water transport (51). In *Aedes* MTs, fluid secretion increases at least three-fold after stimulation with mosquito natriuretic peptide (identified as DH_31_), using cAMP as a second messenger (1), activating PKA, which subsequently activates V-ATPase-driven cation transport processes (22, 35, 39). Herein we show that DH_31_-stimulated secretion is inhibited by bafilomycin, which blocks the proton channel of the VA (32). Moreover, the addition of either bafilomycin or *Aedae*CAPA-1 caused alkalization of the secreted fluid, indicating inhibition of the VA, which may lead to constrained entry of cations across the apical membrane through a proposed alkali cation/proton antiporter (15, 16). Thus, since bafilomycin inhibits DH_31_-stimulated secretion, this supports the VA as a target in the inhibition of fluid secretion.

Consequently, the driving force for ion movement and osmotically-obliged water is reduced, but select Na^+^ channels and cotransporters remain unaffected in the presence of *Aedae*CAPA-1, as observed by the unchanged natriuretic effect of DH_31_ despite reduced secretion rates in response to *Aedae*CAPA-1 (4). Similar results were seen in 5HT-stimulated secretion, albeit a partial inhibition. An earlier study demonstrated that Ca^2+^-mediated diuresis does not require the assembly and activation of the VA (38, 39). The cAMP effect on the VA is implemented by protein kinase A (PKA), with inhibitors of PKA abolishing hormone-induced assembly and activation of the VA (34). Although the endogenous 5HT receptor expressed within the *A. aegypti* MTs necessary for diuretic activity remains elusive, in the kissing bug, *Rhodnius prolixus*, both cAMP and Ca^2+^ have been shown to initiate diuresis in response to 5HT (52), which might explain the partial inhibitory response of *Aedae*CAPA-1 inhibition on 5HT-stimulated tubules as Ca^2+^-mediated diuresis is independent of the VA (39). Notably, the anticipated 5HT type 2 receptor subtype expressed in the principal cells of the MTs is predicted to couple through a Gq/11 signaling mechanism (53) and likely excludes the type 7 Gs-coupled receptor localized to tracheolar cells associated with the MTs (54, 55) as well as the type 1 Gi-coupled receptor localized to principal cells in larval stage mosquitoes (56).

Interestingly, DH_44_-mediated stimulation was observed to be independent of the VA, as bafilomycin had no effect on the secretion rate or pH of the secreted fluid following application of this CRF-related diuretic peptide. Previous studies have noted that low nanomolar concentrations of a DH_44_-related peptide were linked to the stimulation of the paracellular pathway only (27), mediating this action through intracellular Ca^2+^ as a second messenger (57). In contrast, high nanomolar concentrations of a DH_44_-related peptide were shown to influence both paracellular and transcellular transport, increasing intracellular Ca^2+^ and cAMP (57). Although haemolymph concentrations of diuretic peptides have yet to be determined in mosquitoes, DH_31_ is immediately released into circulation post bloodmeal, stimulating rapid secretion of Na^+^ and excess water (23, 50). In contrast, DH_44_-stimulated diuresis in *A. aegypti* involves non-selective transport of Na^+^ and K^+^ cations (4); this supports a delayed release of this diuretic hormone post-feeding to maintain production (albeit reduced) of primary urine whilst conserving Na^+^ ions.

In unstimulated adult female *Aedes* MTs isolated *in vitro*, the VA exhibits variable rates of enzyme activity, consistent with highly variable rates of secretion, as found also in various other insect species (26, 58, 59). The VA is the main energizer in MTs as 60% of total ATPase activity can be linked to the VA (13), whereas the NKA, with around 28% of ATPase activity, also plays a role in membrane energization, denoting a more important role in the function of MTs than was previously assumed (19, 39–41). Here we show a significant two-fold increase in VA activity in MTs treated with DH_31_ compared to the unstimulated MTs, with no change in VA activity in 5HT and DH_44_-treated MTs. Notably, *Aedae*CAPA-1 treatment blocked the DH_31_-driven increase in VA activity, which corroborates the reduced fluid secretion rate and alkalization of the secreted fluid. Additionally, we sought to establish the importance of the NOS/cGMP/PKG pathway in the inhibitory actions of *Aedae*CAPA-1 on VA association. In stimulated MTs treated with *Aedae*CAPA-1 along with NOS inhibitor, _L_-NAME, or PKG inhibitor, KT5823, the inhibitory activity of *Aedae*CAPA-1 and its second messenger cGMP was abolished, resulting in elevated VA activity as a result of DH_31_ treatment. The present study examined the effects of *Aedae*CAPA-1 on DH_31_-stimulated VA activation for the first time in insects. Stimulation of DH_31_ causes an increase in cAMP production, which activates Na^+^ channels and the Na^+^/K^+^/2Cl^-^ cotransporter in the basolateral membrane (60) and up-regulates VA activity (as shown herein and previously) critical for increased fluid secretion (24). The DH_31_ receptor (*Aaeg*GPRCAL1) is expressed in a distal-proximal gradient in the MTs, with greater expression in principal cells where the VA in the apical membrane is highly expressed (19, 61). The co-localization of the DH_31_ receptor, VA, and cation exchangers (62–64) in the distal segment of the MTs, along with the CAPA receptor (6), collectively supports the major roles DH_31_ and CAPA play in post-prandial diuresis and anti-diuresis, respectively. In contrast to the marked changes in VA activity in response to diuretic and anti-diuretic hormones, NKA activity remained unchanged in response to treatments conducted herein. Further studies should examine the potential role of the NKA in diuretic and anti-diuretic processes.

The reversible dissociation of the V_1_ and V_o_ complexes is currently thought of as a universal regulatory mechanism of V-ATPases, appearing to be widely conserved from yeast to animal cells (13, 22, 35, 65). Although previously shown with cAMP (24), it remained unclear whether other second messengers (eg. Ca^2+^, cGMP, and nitric oxide) affect the assembly/disassembly of the V_1_V_o_ complexes in insect MTs. In this study, VA protein abundance in membrane and cytosolic fractions of MTs was confirmed by immunoblot analyses. The 56 kDa band represents the B subunit (29), while the 74 kDa and 32 kDa bands are suggested to be the A and D subunits, respectively, of the V_1_ complex (36). The higher abundance of these V_1_ complex protein subunits in the cytosolic fraction and lower abundance in membrane fraction in *Aedae*CAPA-1-treated MTs provides novel evidence of hormonally-regulated V_1_ dissociation from the holoenzyme in *A. aegypti* MTs. This was further confirmed with V_1_ staining found both in the apical membrane and cytosol of the MTs treated with DH_31_ and *Aedae*CAPA-1 in contrast to the strict co-localization of the V_1_ and V_o_ complex in the apical membrane of MTs treated with DH_31_ alone. This strict co-localization of the VA complex in both DH_31_- and cAMP-treated MTs was further abolished in the presence of cGMP, however, rescued with the addition of the PKG inhibitor, KT5823. In unstimulated *A. aegypti* MTs, 40-73% of the V_1_ subunits were found to be membrane associated, with reassembly of the V_1_V_o_ complex observed upon stimulation with cAMP analogues (39). Although studies have revealed that hormonal regulation can activate the assembly of the holoenzyme, the signaling mechanisms achieving this control are unclear. In this study, the data provides strong evidence of VA assembly in DH_31_-treated MTs, with V_1_ complex protein subunit enrichment found in the membrane fractions, confirming the crucial role of the VA in DH_31_-stimulated secretion. Studies in *A. aegypti* have demonstrated the involvement of PKA in the activation and assembly of the VA upon natriuretic hormone (i.e. DH_31_) stimulation and indicate the phosphorylation of the VA subunits by PKA in the MTs (39). These studies indicate a regulatory role of PKA in VA assembly and its activation that may be independent or in addition to phosphorylation (39). In line with these earlier observations, the current results indicate PKA is critical for DH_31_-stimulated fluid secretion by MTs.

Together, these results indicate that *Aedae*CAPA-1 binds to its cognate receptor in principal cells of the adult *A. aegypti* mosquito MTs (6), targets the NOS/cGMP/PKG pathway (4, 6) to inhibit DH_31_-mediated elevation of cAMP (23, 60), which blocks PKA-activated VA association and prevents protons from being pumped across the apical membrane, resulting in a more alkaline lumen. Additionally, in female *Aedes* MTs, the effect of CAPA peptides are independent of Ca^2+^, which is in contrast to *Drosophila* MTs, where CAPA peptide diuretic activity involves modulation of intracellular concentrations of Ca^2+^, which can derive from both extracellular and intracellular sources of this important signaling molecule (see 9). Our study provides novel evidence that the anti-diuretic activity of CAPA is mediated through the dissociation of the VA holoenzyme involving the removal of the V_1_ complex from the apical membrane, hindering luminal flux of protons that in turn starves cation/H^+^ exchange, which ultimately reduces fluid secretion (**Fig. 6**). CAPA peptides are known to elicit both diuretic and anti-diuretic actions in different insects, whereby a stimulatory role has been established in *D. melanogaster* (9, 10) and inhibitory in the *R. prolixus* (65) and *D. melanogaster* (11, 12) indicating a species-specific role of this neuropeptide family, which may be due to their different diets and lifestyles. Given the unique blood-feeding stress the female mosquito is subjected to, a carefully controlled mechanism of both diuretic and anti-diuretic hormones is warranted to rid the haemolymph of excess salts and water while also preventing the female from excessive secretion.

**Fig 6.**
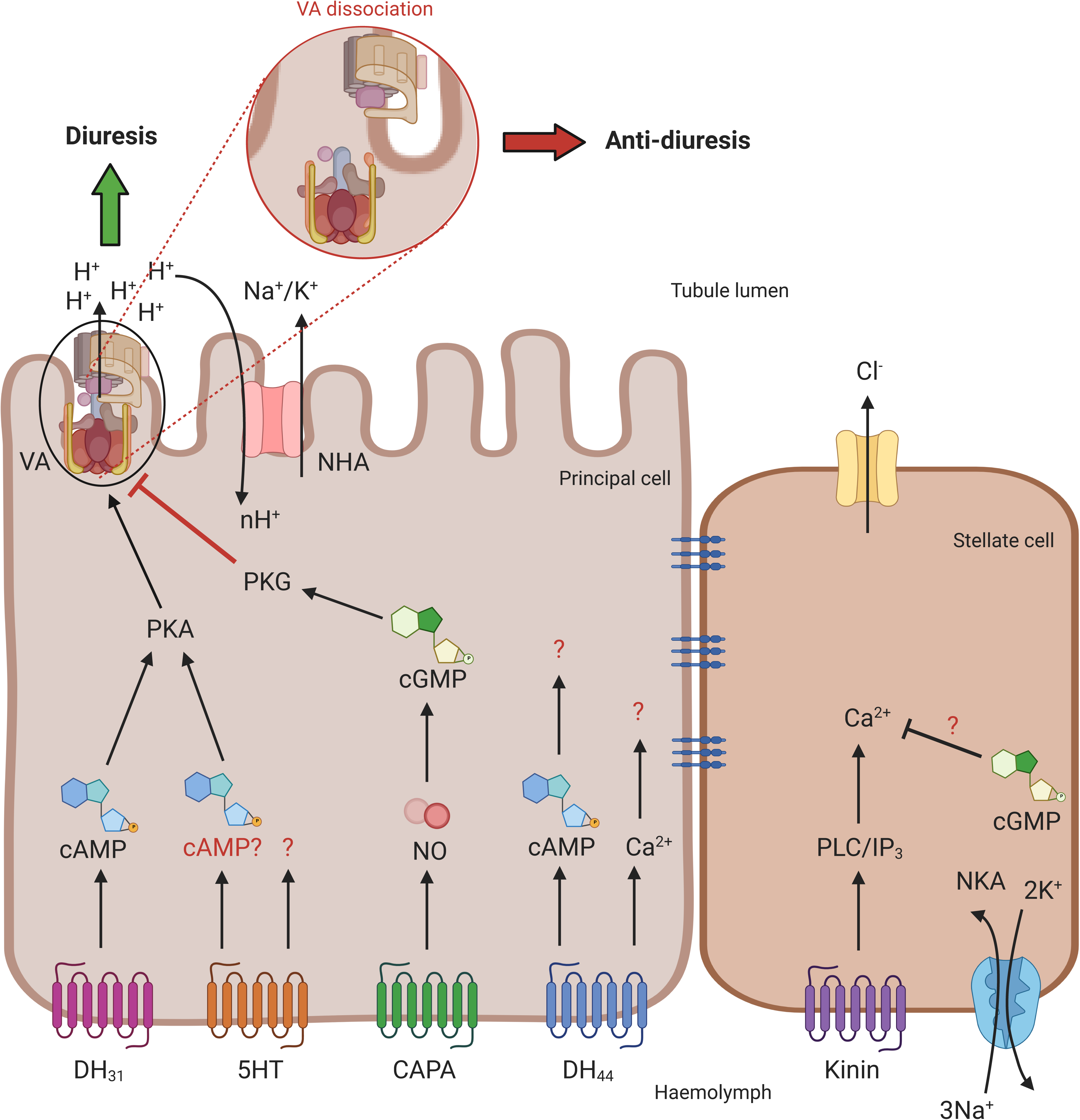
Schematic diagram summarizing the signaling pathway of diuretic and anti-diuretic control of adult *A. aegypti* MTs. The principal cells in *A. aegypti* MTs are responsible for the transport of Na^+^ and K^+^ cations via secondary active transport, energized by the V-type H^+^-ATPase (VA), localized in the brush border of the apical membrane. The movement of protons creates a gradient, driving the exchange of Na^+^ and K^+^ through cation/H^+^ antiporters (NHA). Neurohormone receptors for DH_31_, 5HT, DH_44_, and CAPA are localized to the basolateral membrane of the principal cells, while the kinin receptor is localized exclusively in the stellate cells. The current results together with previous data indicates that DH_31_ stimulates diuresis through activation and assembly of the VA in the apical membrane, with no effect on the Na^+^/K^+^-ATPase (NKA). The anti-diuretic effect of *Aedae*CAPA-1, facilitated by the NOS/cGMP/PKG pathway, causes V_1_ dissociation from the membrane, hindering activity, and thus reducing fluid secretion. The biogenic amine, 5HT, has also been shown to stimulate activation of the VA, however, to a lesser extent. DH_44_-related peptide receptor activation increases Ca^2+^ (as well as cAMP at higher doses) but its action was found to be independent of PKA and VA. Lastly, it was shown earlier that cGMP inhibits kinin-stimulated diuresis, suggesting an additional anti-diuretic factor may exist that acts specifically on stellate cells.

In *R. prolixus* MTs, the physiological roles of cGMP and cAMP were examined (66) suggesting cGMP inhibits fluid secretion by activating a phosphodiesterase (PDE) that degrades cAMP elevated following 5HT and diuretic hormone stimulation of MTs. Indeed, the current results demonstrated the addition of cAMP reversed the inhibitory effects of cGMP, while the addition of cGMP reduced the stimulatory response of cAMP, supporting that these two cyclic nucleotides facilitate two opposing regulatory roles in the MTs of adult *A. aegypti*. The data herein reveals cGMP levels increase in MTs treated with CAPA alone or in combination with DH_31_ while cAMP levels decrease in MTs treated with CAPA in combination with DH_31_ compared to tubules stimulated with DH_31_ alone, which upholds the roles of cAMP and cGMP in diuretic and anti-diuretic signaling pathways, respectively. Interestingly, mid-nanomolar concentrations of DH_44_ also led to increased levels of cAMP, with levels unchanging in response to *Aedae*CAPA-1, raising doubt regarding the involvement of a PDE. While treatment of a PKA inhibitor, KT5720, abolished DH_31_-stimulated secretion, no effect was observed in DH_44_-mediated stimulation. Thus, despite DH_44_-stimulated diuresis was found to involve increased cAMP, it appears to be PKA-independent. It is well established that the effects of cAMP are mediated by activation of cAMP-dependent protein kinase (PKA), a major cAMP target, followed by phosphorylation of target proteins (67). More recently, in *D. melanogaster* MTs, two distinct cAMP pathways have been elucidated to sustain fluid secretion; a PKA-dependent pathway, shown to regulate basal fluid secretion in principal cells; and a PKA-independent pathway, specifically a stimulatory principal EPAC (exchange proteins directly activated by cAMP) pathway, stimulating fluid secretion above basal levels (68). Future studies should examine the DH_44_ diuretic pathway leading to secretion in *A. aegypti* MTs that appears to be PKA-independent and functions in the absence of VA activation.

In summary, our study highlights a novel target in the anti-diuretic signaling pathway of adult female *A. aegypti* MTs, emphasizing the intricate and precise regulatory mechanism of anti-diuresis. Although a plethora studies have investigated the process of hydromineral balance in terrestrial insects from a diuretic perspective (1, 10, 23, 65, 69–71), these current findings advance our understanding of anti-diuretic hormone control while providing further evidence of a previously elusive endocrine regulatory mechanism of the VA in mosquitoes (**Fig 6**). Given that many insects are recognized as agricultural pests or disease vectors, further investigating the complex regulation of their ionic and osmotic balance may aid in lessening their burden on human health and prosperity through development of improved management strategies that, at least in part, impede their neuroendocrine control of hydromineral homeostasis.

## Materials and Methods

### Animal rearing

Eggs of *Aedes aegypti* (Liverpool strain) were collected from an established laboratory colony described previously (4, 72). All mosquitoes were raised under a 12:12 light:dark cycle. Non-blood fed female insects (three-six days post-eclosion) were used for bioassays, dissected under physiological saline (*Aedes* saline) adapted from (60) that contained (in mmol^-1^): 150 NaCl, 25 HEPES, 3.4 KCl, 7.5 NaOH, 1.8 NaHCO_3_, 1 MgSO_4_, 1.7 CaCl_2_, and 5 glucose, and titrated to pH 7.1.

### MT fluid secretion assay

In order to determine fluid secretion rates, modified Ramsay assays were performed as described previously (4, 73). Female adults (3-6 day old) were dissected under physiological *Aedes* saline prepared as described above, and MTs were removed and placed in a Sylgard-lined Petri dish containing 20 μL bathing droplets (1:1 mixture of Schneider’s Insect Medium (Sigma-Aldrich): *Aedes* saline, immersed in hydrated mineral oil to prevent evaporation. The proximal end of each tubule was wrapped around a Minutien pin to allow for fluid secretion measurements. To investigate the effects of second messengers, cAMP and cGMP, on fluid secretion rate, 10^-4^ M 8 bromo-cAMP (cAMP) (23, 66) and 10^-8^ M 8 bromo-cGMP (cGMP) (4) (Sigma-Aldrich, Oakville, ON, Canada) were used against unstimulated MTs. To test the effects of the pharmacological blocker KT5720 (7) (protein kinase A (PKA) inhibitor), a dosage of 5 μmol l^-1^ (manufacturer’s recommended dose) was used against 25 nmol l^-1^ DH_31_- and 10 nmol l^-1^ DH_44_-stimulated MTs. To determine the role of second messenger, Ca^2+^, in the CAPA signaling pathway, MTs were treated in a modified Ca^2+^-free saline (with and without 9.1 mmol^-1^ L-glutamine) containing (in mmol^-1^): 150 NaCl, 25 HEPES, 3.4 KCl, 7.5 NaOH, 1.8 NaHCO_3_, 1 MgSO_4_, and 5 glucose, and titrated to pH 7.1. The role of Ca^2+^ was tested using 1 mmol l^-1^ Ca^2+^ chelator EGTA, 50 μmol l^-1^ (in DMSO) membrane-permeable chelator BAPTA-AM, and 1 nmol l^-1^ intracellular Ca^2+^ chelator 8-(*N*,*N*-diethylamino)octyl 3,4,5-trimethoxybenzoate hydrochloride (TMB-8-HCl) (74).

### Time course inhibition of bafilomycin

Dosage of bafilomycin A_1_ was based on a dose-response analysis of bafilomycin against DH_31_-stimulated tubules (S1 Fig). In the interest of determining whether bafilomycin inhibits the effects of the diuretic factors, dosages of 25 nmol l^-1^ *Drome*DH_31_ (∼84% identical to *Aedae*DH_31_) (4, 23, 75), 100 nmol l^-1^ 5HT (4, 70, 76), and 10 nmol l^-1^ *Rhopr*DH (CRF-related diuretic peptide, DH_44_) (∼48% overall identity; ∼65% identity and ∼92% similarity within the highly-conserved N-terminal region to *Aedae*DH_44_) (4, 57, 77, 78), were applied to the isolated MTs. Neurohormone receptors, including those for 5HT, and the peptides DH_31_, DH_44_, and CAPA, are localized to the basolateral membrane of principal cells (6, 55, 60, 79), while the LK receptor is localized exclusively to stellate cells (5). As a result, the effects of bafilomycin were tested on diuretics known to act on the principal cells of the MTs. After incubating with the individual diuretics for 30 min (using the modified Ramsay assay), diuretic peptide was added alone (controls) or in combination with bafilomycin (final concentration 10^-5^ M). The fluid secretion rate was recorded every 10 min for a total of 80 min. In order to determine whether inhibition of the VA was involved in the anti-diuretic activity of CAPA peptides on adult MTs, the effects of 1 fmol l^-1^ *Aedae*CAPA-1 (4, 6) were investigated in combination with DH_31_ and bafilomycin.

### NKA and VA activities

The Na^+^/K^+^-ATPase (NKA) and VA activity in the MTs was determined using a modified 96-well microplate method (80, 81), which relies on the enzymatic coupling of ouabain- or bafilomycin-sensitive hydrolysis of ATP to the oxidation of reduced nicotinamide adenine dinucleotide (NADH). The microplate spectrophotometer is therefore able to directly measure the disappearance of NADH. Adult female MTs (three to six day old) were dissected and incubated for 30 min in *Aedes* saline, diuretic (DH_31_, 5HT, or DH_44_) alone or combined with *Aedae*CAPA-1. Following 30-min incubation, MTs were collected into 1.5 mL microcentrifuge tubes (40-50 sets of MTs per tube = 200-250 MTs per treatment), flash frozen in liquid nitrogen and stored at -80°C. To investigate the effects of the pharmacological blockers, a nitric oxide synthase (NOS) inhibitor, N_ω_-Nitro-L-arginine methyl ester hydrochloride (_L_-NAME), and protein kinase G (PKG) inhibitor, KT5823, were used against DH_31_-stimulated MTs treated with *Aedae*CAPA-1 or 10^-8^ M cGMP. Dosages of 2 μmol l^-1^ _L_-NAME (manufacturer’s recommended dose) and 5 μmol l^-1^ KT5823 were applied to the MTs (4, 6, 7) (see *SI Materials and Methods for assay preparation*).

### Protein processing and western blot analyses

MTs were isolated under physiological saline from 40-50 female *A. aegypti* for each biological replicate (defined as n = 1) and incubated for 60 min in the following three treatments: *Aedes* saline, 25 nmol l^-1^ DH_31_, or 25 nmol l^-1^ DH_31_ + 1 fmol l^-1^ *Aedae*CAPA-1. Following the incubation, tissues were stored at -80°C until processing. To separate the membrane and cytosolic proteins, a membrane protein extraction kit was used (ThermoFisher Scientific) following recommended guidelines for soft tissue with minor modifications including 200 μL of permeabilization and solubilization buffer and a 1:200 protease inhibitor cocktail (Sigma Aldrich) in both buffers. Final protein concentrations were calculated by Bradford assay (Sigma-Aldrich Canada, Ltd.,) according to manufacturer’s guidelines with bovine serum albumin (BioRad Laboratories) as a standard and quantified using an A_o_ Absorbance Microplate Reader (Azure Biosystems) at 595 nm (see *SI Materials and Methods for western blot analyses*).

### Immunolocalization of VA complexes in MTs

Immunohistochemistry of the MTs localizing the VA complexes was conducted following a previously published protocol (82). Adult female MTs (three to six day old) were dissected out in *Aedes* saline and incubated following similar conditions described in the western blot section above. After the incubation, the MTs were immersed in Bouin’s solution and fixed for two hours in small glass vials. To test *in vivo* changes of the VA complexes, five to six day old females were allowed to blood-feed for 20 min (72), after which female mosquitoes were isolated at 10 min, 30 min, and 3 hr post blood-meal. Similarly aged, non blood-fed (sucrose-fed) females were isolated as controls. Following the bloodmeal, whole body females were immersed in Bouin’s solution and fixed for three hours in small glass vials. To test role of second messengers in VA localization, similarly aged female MTs were incubated in solutions containing 10^-4^ M cAMP, 10^-8^ M cGMP, and 5 μmol l^-1^ KT5823 before being immersed in Bouin’s solution and fixed for two hours. Tissues/whole body females were then rinsed three times and stored in 70% ethanol at 4°C until further processing. Fixed samples were dehydrated through a series of ethanol washes: 70% ethanol for 30 min, 95% ethanol for 30 min, and 100% ethanol three times for 30 min. The samples were cleared with xylene (ethanol:xylene for 30 min then 100% xylene three times for 30 min), and infiltrated in Paraplast Plus Tissue Embedding Medium (Oxford Worldwide, LLC, Memphis, USA) at xylene:paraffin wax for 60 min at 60°C, then rinsed in pure paraffin wax twice for 1 hour for 60°C. Following the last rinse, the samples were embedded in the paraffin wax and left to solidify at 4°C until further processing. Sections (5 μm) were cut using a Leica RM 2125RT manual rotary microtome (Leica Microsystems Inc., Richmond Hill, Canada) and slides were incubated overnight on a slide warmer at 45°C for preparation of immunohistochemistry (see *SI Materials and Methods*).

### cGMP and cAMP Measurements

A competitive cGMP ELISA kit (Cell Signaling Technology, #4360) and cAMP ELISA kit (Cell Signaling Technology, #4339) were used to measure these cyclic nucleotide second messengers from adult *A. aegypti* MTs following different treatments (see *SI Materials and Methods for further details*).

### Statistical analyses

Data was compiled using Microsoft Excel and transferred to Graphpad Prism software v.7 to create figures and conduct all statistical analyses. Data was analyzed accordingly using a one-way or two-way ANOVA and a Bonferroni post-test, or Student’s t-test, with differences between treatments considered significant if p<0.05.

## Author Contributions

J.-P.P. and F.S. designed research; F.S. and M.F.V. performed research; F.S. and J.P.P. analyzed data; and J.-P.P. and F.S. wrote and revised the manuscript.

## Classification

Biological Sciences; Physiology

## Acknowledgments

The authors sincerely thank Prof. Helmut Wieczorek and Prof. Felix Tiburcy (University of Osnabrück, Germany) for providing the V_1_ antibody and Prof. Andrew Donini (York University, Canada) for providing the *A. aegypti* AQP1 antibody that were used in this study. The authors are also grateful to Prof. Ian Orchard (University of Toronto Mississauga, Canada) and Prof. Michael J. O’Donnell (McMaster University, Canada) for providing synthetic peptides, *Rhopr*DH and *Drome*DH_31_ respectively, used in this study. The authors would like to thank the reviewers for their insightful comments and feedback which helped improve an earlier version of this manuscript.

## Data and materials availability

All data are available in the main text of the article or the supplementary information.

