## Supplementary information for "The V-type H^+^-ATPase is targeted in anti-diuretic hormone control of the Malpighian ‘renal’ tubules"

**Author Contributions:** J.-P.P. and F.S. designed research; F.S. and M.F.V. performed research; F.S. and J.P.P. analyzed data; and J.-P.P. and F.S. wrote and revised the manuscript.
**Competing Interest Statement:** The authors declare no competing interest.
**Classification:** Biological Sciences; Physiology
**Keywords:** excretory system, *Aedes aegypti*, mosquito, anti-diuresis, proton pump, disease vector

**This PDF file includes:**

Supplementary Text
Figures S1 to S7

### SI Materials and Methods

#### NKA and VA activities

Tissues were thawed on ice and 150 μL of homogenizing buffer (four parts (in mmol $l^{-1}$) of: 150 sucrose, 10 EDTA, and 50 imidazole (SEI); pH 7.3 and one part of SEID with composition: 0.5% of sodium deoxycholic acid in SEI) and MTs were then sonicated on ice for 10 sec (two pulses of 5 sec) and subsequently centrifuged at 10,000 x *g* for 10 min at 4°C. The supernatant was transferred into a fresh microcentrifuge tube and stored on ice.

Prior to the assay, three solutions were prepared (Solution A, B, and C) and all stored on ice. Solution A contained 4 units $mL^{-1}$ lactate dehydrogenase (LDH), 5 units $mL^{-1}$ pyruvate kinase (PK), 50 mmol $l^{-1}$ imidazole, 2.8 mmol $l^{-1}$ phosphoenolpyruvate (PEP), 0.22 mmol $l^{-1}$ ATP, and 50 mmol $l^{-1}$ NADH, pH 7.5. The solution was subsequently mixed with a salt solution in a 4:1 ratio (salt solution composition (in mmol $l^{-1}$): 189 NaCl, 10.5 $MgCl_2$, 42 KCl, and 50 imidazole, pH 7.5. Working solution B consisted of solution A with 5 mmol $l^{-1}$ ouabain and solution C consisted of solution A with 10 μmol $l^{-1}$ bafilomycin. The concentrations of ouabain and bafilomycin were based on previous studies (1). To ensure the batch of assay mixture (Solution A) was effective, an adenosine diphosphate (ADP) standard curve was run. ADP standards were prepared as follows: 0 nmol 10μ$L^{-1}$ (200μL of 50mmol $l^{-1}$ imidazole buffer (IB), pH 7.5); 5 nmol 10μ$L^{-1}$ (25μL of 4mmol $l^{-1}$ ADP stock and 175μL of IB); 10 nmol 10u$L^{-1}$ (50μL of 4mmol $l^{-1}$ ADP stock/ 150 μL of IB); 20 nmol 10 μ$L^{-1}$ (100μL of 4mmol $l^{-1}$ ADP stock/100μL of IB); 40 nmol 10 μ$L^{-1}$ (40 mmol $l^{-1}$ ADP stock). The standards were added to a 96-well polystyrene microplate in duplicates of 10 μL per well, followed by the addition of 200 μL of solution A. The plate was placed in a Thermo Multiscan Spectrum microplate spectrophotometer set at 25°C and a linear rate of NADH disappearance was measured at 340 nm. The absorbance spectra were recorded and analyzed using the Multiscan Spectrum data acquisition system with SkanIt version 2.2 software. The ADP standards (0 to 40 nmoles well$^{-1}$) should yield an optical density (OD) between 0.9 and 0.2 OD units, while the slope of the curve should result in -0.012 to -0.014 OD nmol ADP$^{-1}$. Homogenized MT samples were thawed and added to the microplate (kept on ice) in six replicates of 10 μL per well. Next, two wells per sample were filled with 200 μL of working solution A, two wells with 200 μL of working solution B and two wells with 200 μL of working solution C. The microplate was quickly placed in the microplate spectrophotometer and the decrease in NADH absorbance was measured for 30 min at 340 nm. NKA and VA activity was calculated using the following equation:

$$\text{NKA or VA activity} = (((\Delta\text{ATPase}/S)/[\text{P}]) \times 60 \text{ (min)},$$

where ΔATPase is the difference in ATP hydrolysis in the absence and presence of ouabain or bafilomycin, *S* is the slope of the ADP standard curve, [P] is the protein concentration of the sample. Protein was quantified using a Bradford assay (Sigma-Aldrich Canada, Ltd.,) according to manufacturer's guidelines with bovine serum albumin (Bioshop Canada Inc., Burlington, ON, Canada) as a standard. Final activity was expressed as micromoles of ADP per milligram of protein per hour.

**Measurement of pH of the secreted fluid**

The pH of secreted fluid was measured by using ion-selective microelectrodes (ISME) pulled from glass capillaries (TW-150-4, World Precision Instruments, Sarasota, Fl, USA) using a Sutter P-97 Flaming Brown pipette puller (Sutter Instruments, San Raffael, CA, USA). Microelectrodes were silanized with *N,N*-dimethyltrimethylsilylamine (Fluka, Buchs, Switzerland) pipetted onto the interior of a glass dish inverted over the group of microelectrodes. A 1:2 ratio of number of microelectrodes to amount of silanization solution (in µl) was used. The microelectrodes were left to silanize for 75 min at 350°C and left to cool before use. The microelectrodes were back-filled with a solution containing 100 mmol $l^{-1}$ NaCl and 100 mmol $l^{-1}$ sodium citrate that was titrated to pH 6.0 and front-filled using Hydrogen Ionophore I – cocktail B (Fluka, Buchs, Switzerland). The electrode tips were then coated with ~3.5% (w/v) polyvinyl chloride (PVC) dissolved in tetrahydrofuran, to avoid displacement of the ionophore cocktail when submerged in the paraffin oil (2). The $H^+$-selective microelectrodes were calibrated in *Aedes* saline titrated to either pH 7.0 or pH 8.0. Reference electrodes were prepared from glass capillaries (1B100F-4, World Precision Instruments) using a pipette puller described above and were backfilled with 500 mmol $l^{-1}$ KCl. Secreted droplet pH measurements were done immediately after collection to prevent alkalization of the droplet due to carbon dioxide diffusion into the paraffin oil. Microelectrodes and reference electrodes were connected to an electrometer through silver chloride wires where voltage signals were recorded through a data acquisition system (Picolog for Windows, version 5.25.3). In order to measure pH of the secreted fluid, tubules were set up using the Ramsay assay, and pH measurements were recorded every 10 min for a total of 60 min.

**cGMP and cAMP Measurements**

A competitive cGMP ELISA kit (Cell Signaling Technology, #4360) and cAMP ELISA kit (Cell Signaling Technology, #4339) were used to measure the effect of $DH_{31}$, $DH_{44}$ and *Aedae*CAPA-1 on cGMP and cAMP levels in the MTs. Adult MTs were isolated under physiological saline from 50 female *A. aegypti* for each biological replicate (defined as n = 1). To prevent cGMP degradation, tubules were incubated first with a phosphodiesterase inhibitor, 0.1 mmol $l^{-1}$ zaprinast, for 10 min (3) or 0.5 mmol $l^{-1}$ 3-isobutyl-1-methylxanthine (IBMX) for 10 min (4) before any of the experimental treatments, specifically *Aedes* saline, 25 $nmol^{-1}$ $DH_{31}$, 10 nmol $l^{-1}$ $DH_{44}$, 1 $fmol^{-1}$ *Aedae*CAPA-1, 25 $nmol^{-1}$ $DH_{31}$ + 1 $fmol^{-1}$ *Aedae*CAPA-1, or 10 $nmol^{-1}$ $DH_{44}$ + 1 $fmol^{-1}$ *Aedae*CAPA-1 for a further 20 min. After incubation was complete, tissues were stored at -80ºC until processing. To measure cGMP and cAMP concentrations, frozen tubule samples were thawed on ice and 125 µL of 1X cell lysis buffer (CLB, #9803) was added to each tube (1 $mmol^{-1}$ phenylmethylsulfonyl fluoride (PMSF) was added to 1X CLB fresh each time). Tissue samples were kept on ice for 10 min, sonicated for 10 s (similar conditions as described for NKA/VA activity assay), centrifuged for 3 min at 10,000 rpm, and the supernatant was isolated and kept on ice. Using a commercial 96-well microtitre plate precoated with either a cGMP or cAMP rabbit monoclonal antibody, the cGMP-HRP (or cAMP-HRP) conjugate was added in triplicate wells with 50 µL per well. This was followed by the addition of 50 µL of either tubule samples or cGMP (or cAMP) standards ranging from 100 $nmol^{-1}$ to 0.25 $nmol^{-1}$. The plate was covered and incubated at RT for 3 h on a horizontal orbital plate shaker. Following incubation, plate contents were discarded, and wells were washed four times with 200 μL/well of 1X wash buffer. Next, 100 μL of 3,3',5,5'-tetramethylbenzidine (TMB) substrate was added to each well, and the plate was covered and kept for 10 min at RT. The enzymatic reaction was quenched by adding 100 μL of 2 $mmol^{-1}$ HCl and absorbance read at 450 nm using a Synergy 2 Microplate Reader (Biotek).

**Western blot analyses**

Protein samples were denatured by heating for 5 min at 100ºC with 6X loading buffer (225 mmol $l^{-1}$ Tris-HCl pH 6.8, 3.5% (w/v) SDS, 35% glycerol, 12.5% (v/v) ß-mercaptoethanol and 0.01% (w/v) Bromophenol blue). Into each lane, 5 μg of protein was loaded onto a 4% stacking and 12% resolving sodium dodecyl sulphate polyacrylamide gel electrophoresis (SDS-PAGE) gel. Protein samples were migrated initially at 80 V for 30 min and subsequently at 110 V for 90 min before being transferred onto a polyvinylidene difluoride (PVDF) membrane using a wet transfer method at 100 V for 60 min in a cold transfer buffer. Following transfer, PVDF membranes were blocked with 5% skim milk powder in Tris-buffered saline (TBS-T; 9.9 mmol $l^{-1}$ Tris, 0.15 mmol $l^{-1}$ NaCl, 0.1 mmol $l^{-1}$ Tween-20, 0.1 mmol $l^{-1}$ NP-40 pH 7.4) for 60 min at RT, and incubated on a rocking platform overnight at 4ºC with a guinea pig polyclonal anti-VA (Ab 353-2 against the $V_1$ complex of the VA (5), a kind gift from Profs. Wieczorek and Tiburcy, University of Osnabruck, Germany, used at a 1:2000 dilution in 5% skim milk in TBS-T. The next day, PVDF membranes were washed for 60 min in TBST-T, changing the wash buffer every 15 min. Immunoblots were then incubated with a goat anti-guinea pig HRP conjugated secondary antibody (1:2500 in 5% skim milk in TBS-T)

(Life Technologies, Burlington, ON) for 60 min at RT and subsequently washed three times for 15 min with TBS-T. Lastly, blots were incubated with the Clarity Western ECL substrate and images were acquired using a ChemiDoc MP Imaging System (Bio-Rad). Molecular weight measurements were performed using Image Lab 5.0 software (Bio-Rad). PVDF membranes were then probed with Coomassie brilliant blue, since total protein normalization is now considered the benchmark method for quantitative analysis of western blot data (6, 7) and has been used in studies involving *A. aegypti* protein normalization (8). ImageJ software (NIH, USA) was used to quantify protein abundance. (**Fig 3** and **S5 Fig** show a saturated blot of the 56kDa band to ensure visualization of the 74kDa and 32kDa band, however protein quantification was measured using a pre-saturated blot).

To confirm successful separation of membranes from cytosol using the membrane protein extraction kit, saline-incubated MTs were incubated in either anti-beta-tubulin (cytosolic marker, 1:5000) or -AaAQP1 affinity purified rabbit polyclonal antibody (generous gift from Dr. Andrew Donini, York University, Canada) (9) (membrane marker, 1:1000). Blots were then incubated with a goat anti-mouse (for beta-tubulin) or goat anti-rabbit (for AQP1) HRP-conjugated secondary antibody (1:5000 in 5% skim milk in TBS-T) (Life Technologies, Burlington, ON) for 60 min at RT.

**Immunolocalization of VA complexes in MTs**

Tissue sections were deparaffinized with xylene (two rinses for 5 min each), and rehydrated via a descending series of ethanol washes (100% ethanol twice for 2 min, 95% ethanol for 2 min, 70% ethanol for 2 min, 50% ethanol for 2 min) and finally in distilled water for 20 min. Next, sections were subjected to a heat-induced epitope retrieval (HIER) by immersing slides in a sodium citrate buffer (10 nmol $l^{-1}$, pH 6.0) and heating both slides and solution in a microwave oven for 4 min. The solution and slides were allowed to cool for 20 min, reheated for 2 min, and left to stand at room temperature (RT) for 15 min. Slides were then washed three times in phosphate-buffered saline (PBS) pH 7.4, 0.4% Kodak Photo-Flo 200 in PBS (PBS/PF, 10 min), 0.05% Triton X-100 in PBS (PBS/TX, 10 min), and 10% antibody dilution buffer (ADB; 10% goat serum, 3% BSA and 0. 05% Triton X-100 in PBS) in PBS (PBS/ADB, 10 min). Slides were incubated overnight at RT with a guinea pig polyclonal anti-$V_1$ (Ab 353-2 against the $V_1$ complex, identical antibody used in western blot analyses, at 1:5000 dilution in ADB) in combination with a 1:100 mouse polyclonal anti-ATP6V0A1 antibody for $V_o$ (Abnova, Taipei, Taiwan).

Following overnight primary antibody incubation, slides were washed briefly in distilled water, with sequential washes with PBS/PF, PBS/TX, and PBS/ADB for 10 min each as described above. A goat anti-guinea pig antibody (for $V_1$ detection) conjugated to AlexaFluor 488 (1:500 in ADB, Jackson Immunoresearch) and sheep anti-mouse antibody (for $V_0$ detection) conjugated to AlexaFluor 594 (1:500 in ADB, Jackson Immunoresearch) were applied to visualize the VA complexes. The slides were left to incubate in the secondary antibody for 60 min at RT. For negative controls, slides were processed as described above with primary antibodies omitted. Slides were then washed in distilled water/0.4% PF three times for 1 min each and finally in distilled water for 1 min. Slides were air dried for 30 min and mounted using ProLong$^{TM}$ Gold antifade reagent with DAPI (Life Technologies, Burlington, ON, Canada). Fluorescence images were captured using an Olympus IX81 inverted fluorescent microscope (Olympus Canada, Richmond Hill, ON, Canada).

### Supplemental Information References

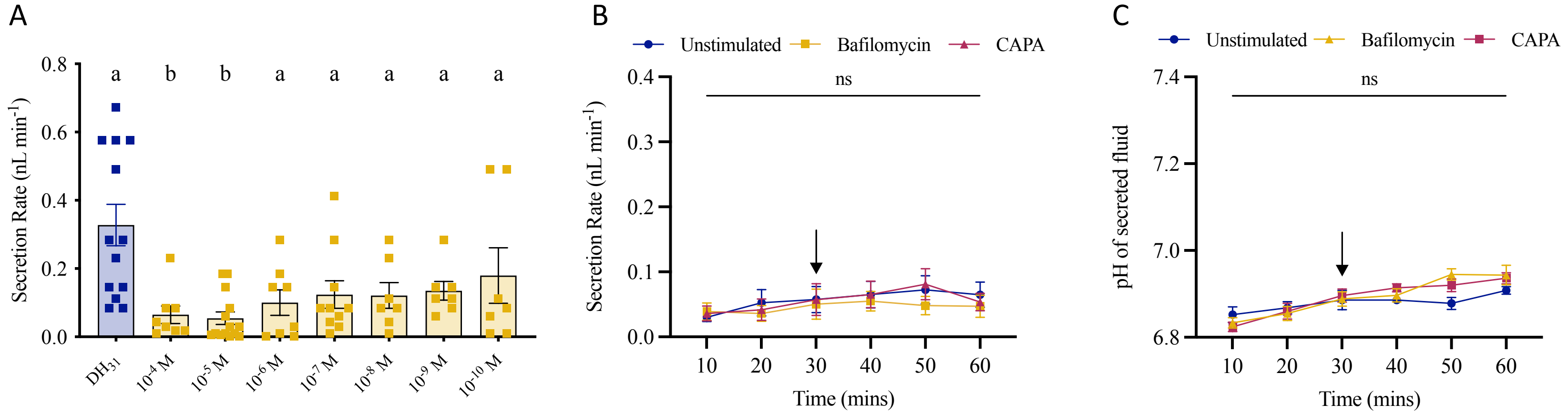


**S1 Fig. Dose response and effect of bafilomycin on fluid secretion rates and pH of MTs *in vitro* isolated from adult female *A. aegypti.***

(A) Doses of $10^{-4}$ M to $10^{-10}$ M bafilomycin were applied to MTs together with $DH_{31}$ for 60 min. Bars labeled with different letters are significantly different from each other (mean± SEM; one-way ANOVA with Bonferroni multiple comparison, $p<0.05$, n = 7–13). (B) Fluid secretion rates were measured at 10 min intervals over a 30 min control period and then after the addition (solid arrow) of $10^{-5}$ M bafilomycin (yellow), 1 fmol $l^{-1}$ *Aedae*CAPA-1 (purple), or unstimulated alone (blue). (C) The pH of the secreted droplets was measured at 10-min intervals for 60 min using an ion-selective microelectrode. No significant differences were observed in the measurements (mean± SEM; two-way ANOVA with Bonferroni multiple comparison, $p<0.05$, n = 5–8); ns denotes no statistical significance.

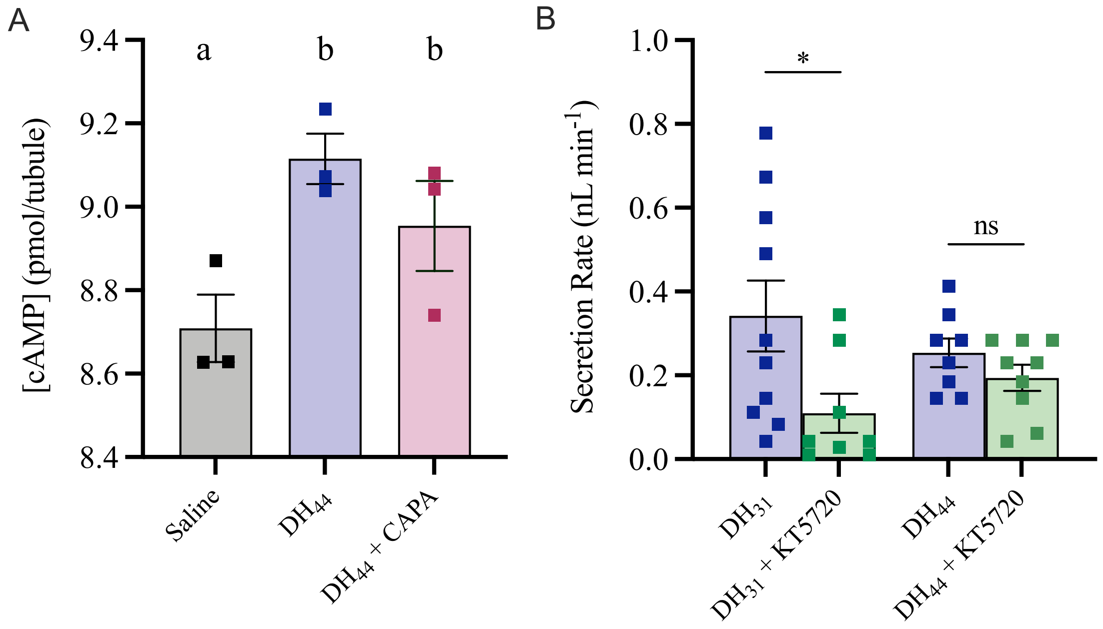


**S2 Fig. Intracellular levels of cAMP in $DH_{44}$-treated *A. aegypti* MTs.**
(A) To monitor activity of *Aedae*CAPA-1 on cAMP levels in $DH_{44}$-stimulated *A. aegypti* MTs, tubules were incubated in IBMX for 10 min prior to the following treatments; *Aedes* saline, $DH_{44}$ alone, or in combination with *Aedae*CAPA-1 for 20 min. Bars labeled with different letters are significantly different from each other (mean± SEM; one-way ANOVA with Bonferroni multiple comparison, $p<0.05$. For each treatment, 50 sets of MTs were isolated and combined (n = 3 biological replicates per treatment). (B) Female MTs were treated with either $DH_{31}$ or $DH_{44}$ alone or in combination with the PKA inhibitor, KT5720 for 60 min to measure fluid secretion rate. Significant differences are denoted by an asterisk (mean± SEM; one-way ANOVA with Bonferroni multiple comparison, $p<0.05$, n = 8–10).

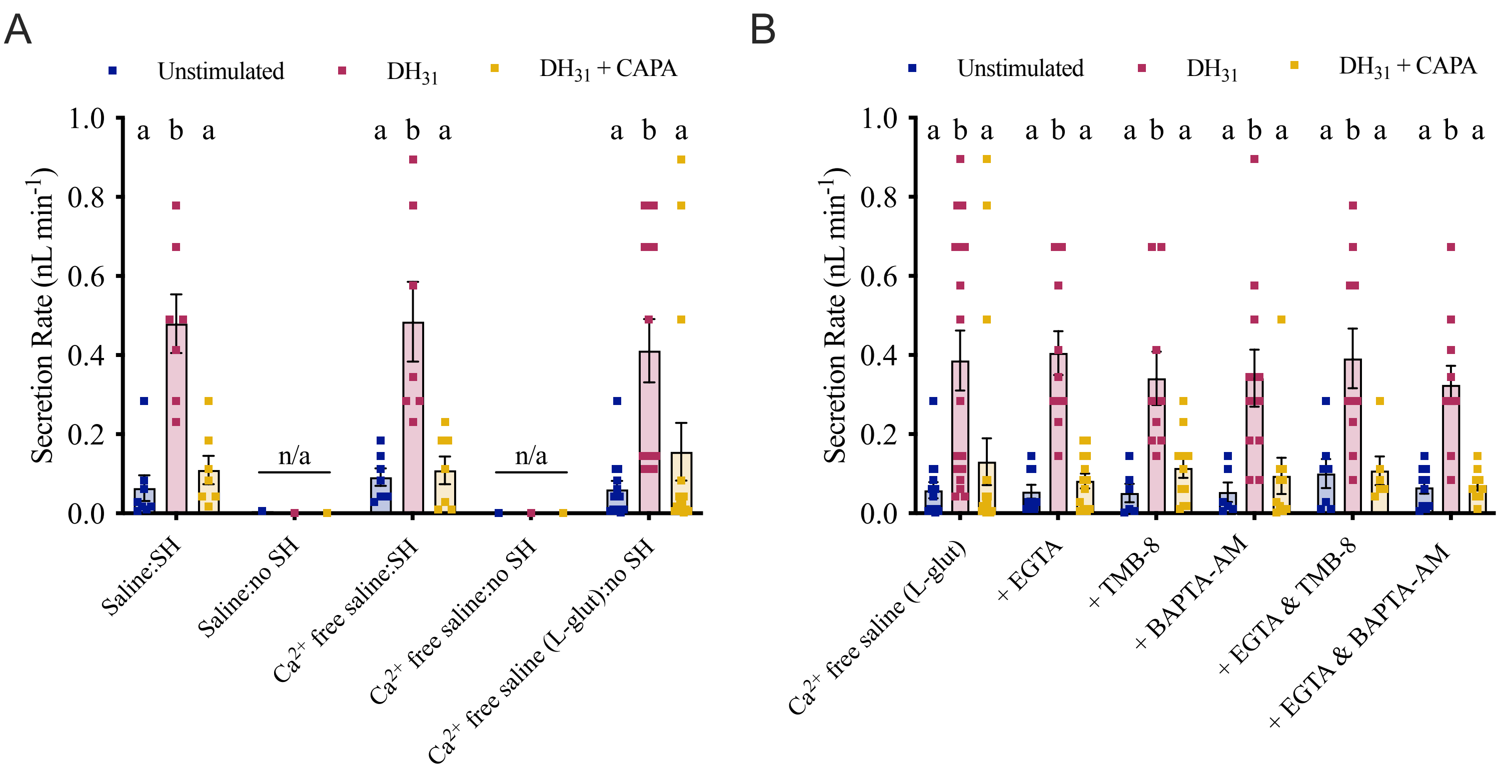


**S3 Fig. Effect of $Ca^{2+}$-free saline, EGTA, TMB-8, and BAPTA-AM on fluid secretion rate in $DH_{31}$- and CAPA-treated *A. aegypti* MTs.**

(A) MTs were treated with $DH_{31}$ alone or in combination with *Aedae*CAPA-1 in solutions containing; *Aedes* saline with/without Schneider's medium (SH), $Ca^{2+}$-free *Aedes* saline with/without Schneider's medium (SH), and a modified $Ca^{2+}$-free saline with L-glutamine (see *Methods* section in main manuscript). Fluid secretion rate was measured after 60 min. (B) Female MTs were incubated in a modified $Ca^{2+}$-free saline (L-glutamine) with $DH_{31}$ alone or in combination with *Aedae*CAPA-1 in the presence of blockers, EGTA, TMB-8, and/or BAPTA-AM for 60 min. Bars labeled with different letters are significantly different from each other (mean± SEM; one-way ANOVA with Bonferroni multiple comparison, $p<0.05$, n = 6–20, n/a = not applicable).

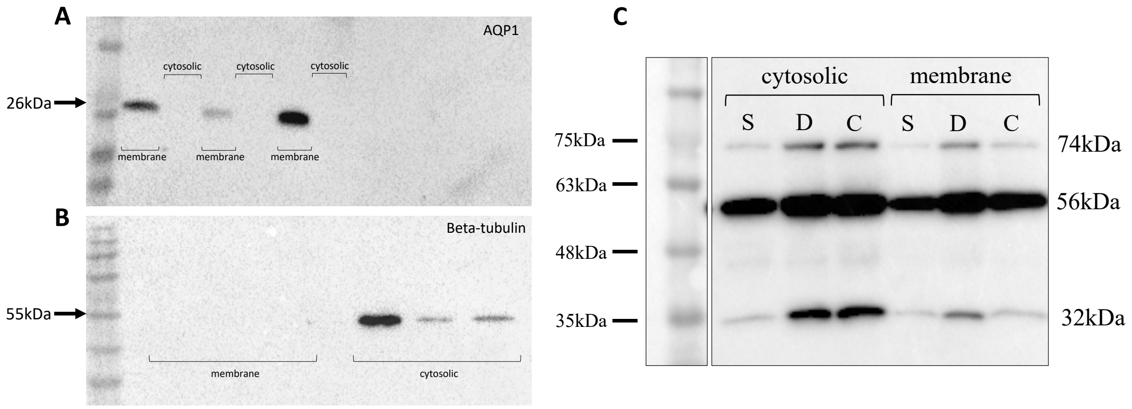


**S4 Fig. Representative western blot of membrane and cytosolic protein fractions.** (A) Membrane and cytosolic protein fraction isolation was validated using established membrane (aquaporin-1 in *A. aegypti* MTs) and cytosolic (beta-tubulin) protein markers. Aquaporin-1 (AQP1) detection shown in membrane fractions of *Aedes* saline-incubated MTs at 26 kDa, and (B) beta-tubulin in cytosolic fractions of *Aedes* saline-incubated MTs at 55 kDa (n=40–50 MTs per replicate, n = 3 biological replicates). (C) The MTs (n = 40–50) were incubated in *Aedes* saline alone, $DH_{31}$, or $DH_{31}$ + *Aedae*CAPA-1 for one hour before collection. Western blot analysis revealed three protein bands, with calculated molecular masses of 74 kDa, 56 kDa, and 32 kDa. In panel B, S = *Aedes* saline, D = $DH_{31}$, and C = $DH_{31}$ + *Aedae*CAPA-1.

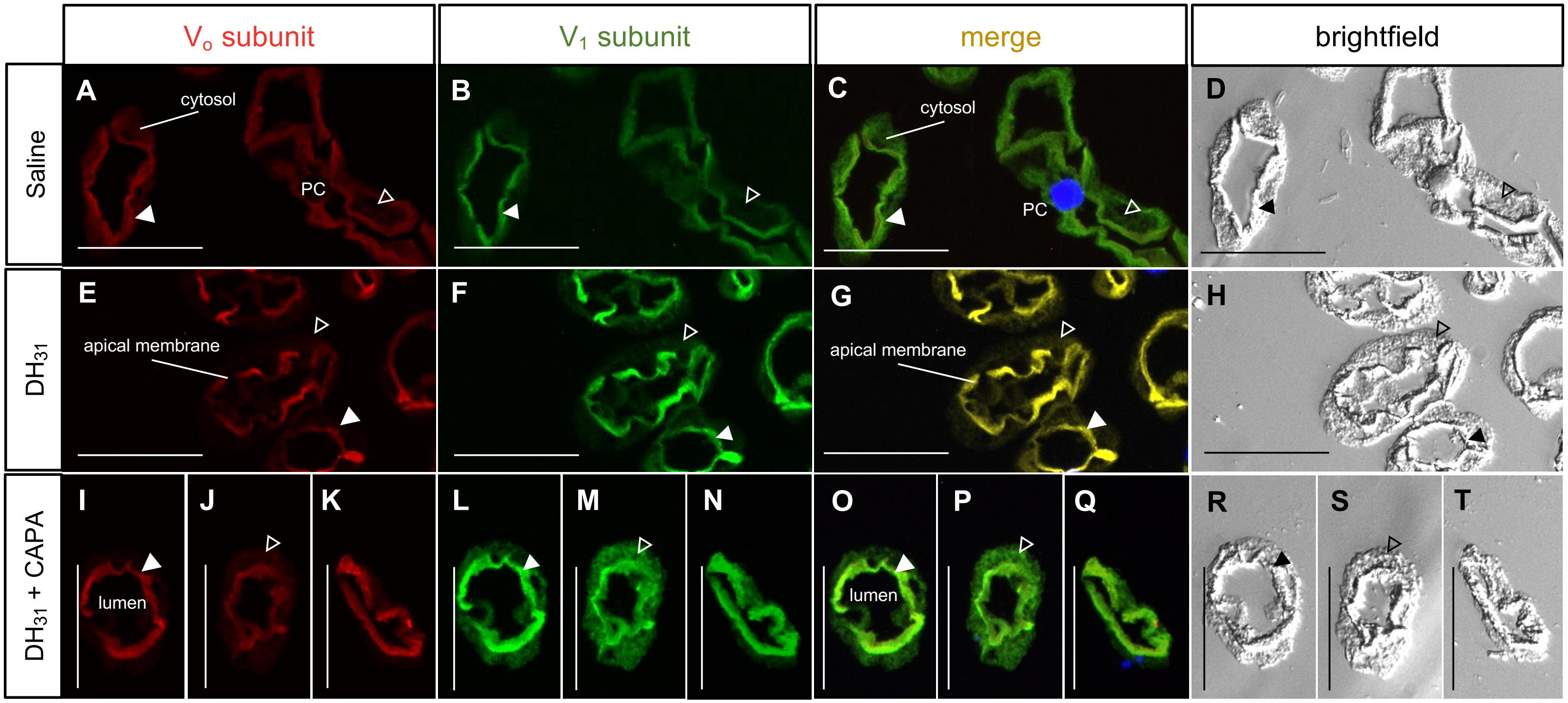


**S5 Fig. Immunolocalization of the $V_o$ and $V_1$ complexes in cross sections of stimulated *A. aegypti* MTs.**
Representative paraffin-embedded sections of *A. aegypti* MTs incubated in either (A–D) *Aedes* saline, (E–H) $DH_{31}$ and (I–T) $DH_{31}$ + *Aedae*CAPA-1. Panels (A,E,I–K) show membrane-integrated $V_o$ immunoreactive staining (red), (B,F,L-N) demonstrate $V_1$ immunoreactive staining (green), (C,G,O–Q) show merged images of $V_o$ and $V_1$ staining, while (D,H,R–T) show brightfield images. Solid white arrowheads denote apical VA staining, and empty arrowheads mark cytosolic VA staining. DAPI nuclear staining is shown in blue. Scale bar 100 µm, n = 3–4 biological replicates (PC = principal cell).

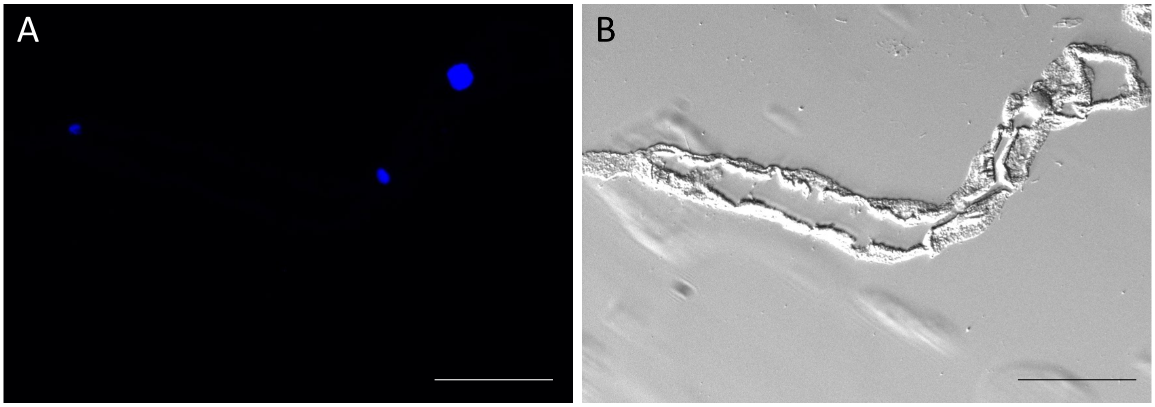


**S6 Fig. Negative controls of $V_0$ and $V_1$ complexes in cross sections of *A. aegypti* MTs.** Paraffin-embedded sections incubated in no primary controls. (A) No staining was observed in the MTs while (B) show representative brightfield image. DAPI nuclear staining is shown in blue. Scale bar 100 μm.

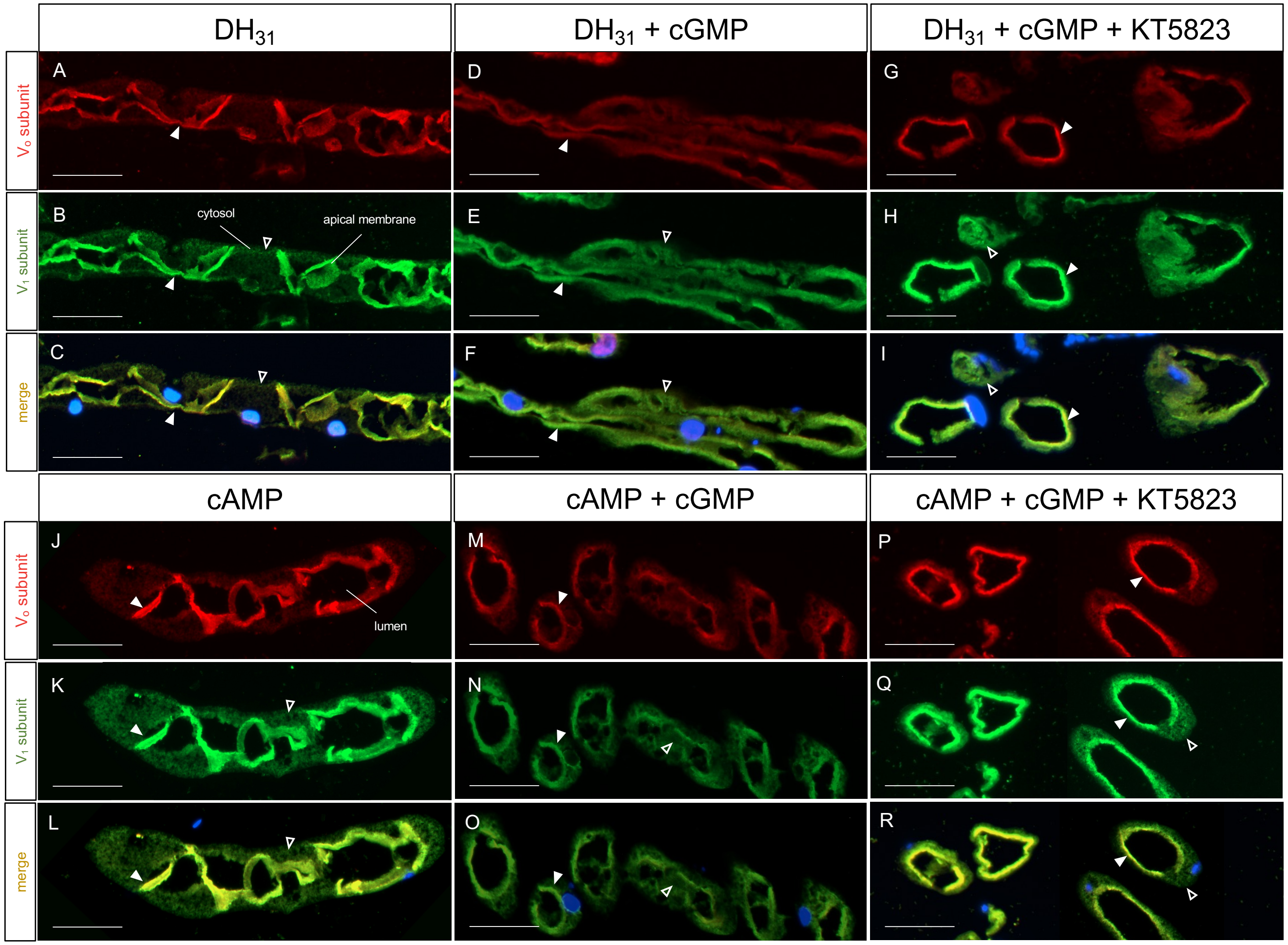


**S7 Fig. Effect of second messengers, cAMP and cGMP, and PKG inhibition on immunolocalization of $V_o$ and $V_1$ complexes in transverse sections of *A. aegypti* females.**

Representative paraffin-embedded sections of *A. aegypti* MTs incubated in either (A–C) $DH_{31}$ alone, $DH_{31}$ combined with cGMP (D–F), or $DH_{31}$ combined with cGMP + KT5823 (G–I). Similar representative paraffin-embedded sections of tubules treated with (J–L) cAMP alone, cAMP combined with cGMP (M–O), or cAMP combined with cGMP + KT5823 (P–R). Panels (A,D,G,J,M,P) show membrane-integrated $V_o$ immunoreactive staining (red), (B,E,H,K,N,Q) demonstrate $V_1$ immunoreactive staining (green), and (C,F,I,L,O,R) show merged images of $V_o$ and $V_1$ staining. Solid white arrowheads denote apical VA staining, and empty arrowheads mark cytosolic VA staining. DAPI nuclear staining is shown in blue. Scale bar 100 µm, n = 3 biological replicates.
